# Community resilience in a river network: the roles of connectivity and drying regime

**DOI:** 10.1101/2025.07.11.664354

**Authors:** Amélie Truchy, Romain Sarremejane, Eléonore Braun, Thibault Datry

## Abstract

The cross-scale resilience model suggests that resilience, the amount of disturbance an ecosystem can absorb before collapsing and reorganizing, can be measured by evaluating the diversity and redundancy of functions performed by species at different spatiotemporal scales. Yet, little is known about the effects of flow intermittence and associated hydrological connectivity on the resilience capacity of instream communities, and the ecosystem functions they perform. We expected lower resilience capacity in non-perennial and isolated reaches. Here, we used fish and invertebrate community data and litter decomposition rates across 20 sites in a river network naturally fragmented by drying to characterize the drivers of resilience at the river-network scale. Using discontinuity analysis, a set of resilience indicators were calculated from body size distribution and species traits, and related to flow intermittence, network fragmentation and position in the stream network. We found that non-perennial reaches were characterized by lower resilience with fewer species, lower levels of functional redundancy of five out of eight functional feeding groups. Functional redundancy and response diversity in shredders were decoupled, translating into low litter decomposition rates in non-perennial reaches. Upstream reaches were characterized by low community resilience, likely reflecting their isolated position in the river network, but flow intermittence affected more strongly the resilience of downstream than upstream reaches. Cross-scale functional redundancy and grazer response diversity were driven by network fragmentation, meaning that the functions these groups perform might be at risk when facing other anthropogenic pressures. Finally our study suggests that reliable resilience assessments need to be based on several standardized indicators and call for more studies comparing these indicators in diverse ecosystems.

## Introduction

Although regional and local community diversity are increasingly threatened by global change, a single facet of biodiversity – species richness – has historically been the overwhelming metric used to analyse and predict changes in ecosystem functioning and resilience (Oliver et al. 2015, Vellend 2017, Ingrisch & Bahn 2018). However, the relationship between biodiversity and ecosystem functioning is underpinned by the diversity of functional roles performed by the species in a community (e.g. Tilman et al. 2001, Hooper et al. 2002). For example, an increase in species richness might have marginal effects on ecosystem functioning if new colonizers are functionally redundant with some species already present in the community (Loreau et al. 2002, Elmqvist et al. 2003, Biggs et al. 2020). Inversely, an increase in the dominance of a single species that plays a unique functional role might have disproportionate effects on functioning, without a change in species richness (McKie et al. 2008, Crawford et al. 2021, Eisenhauer et al. 2023). Such mismatches in functional and structural responses highlight the importance of combining both facets, in understanding and regulating ecosystem functioning and resilience, particularly in the context of current global changes (Truchy et al. 2015, Crabot et al. 2021, Biggs et al. 2020).

Resilience theory aims at predicting complex nonlinear shifts in communities and ecosystems. Ecological resilience is the amount of change needed to force a system from being maintained by one set of processes and structures to a different system regulated by another set of processes and structures (Holling 1973, Angeler & Allen 2016, Van Looy et al. 2019). Once disturbance thresholds are crossed, ecosystems can reorganize in alternative states, (i.e. regime shifts; Beisner et al. 2003) with new sets of processes and structures reinforcing the new state and preventing transition between alternative states (Smol et al. 2007, Dakos et al. 2019). Ecological resilience is related to – but distinct from – ecological stability, also called “engineering resilience”, “resistance”, “elasticity” and “bounce-back time” (Angeler et al. 2016, Hillebrand et al. 2018, Gianuca et al. 2024). The distinction is crucial as ecological resilience looks at regime shifts between alternative states, while ecological stability considers only a single regime (Angeler & Allen, 2016). Therefore, quantifying ecological resilience should be done distinctly from stability and a number of metrics has been flourishing in the research field of ecology to do so (Allen et al. 2005, Baho et al. 2017).

The cross-scale resilience model, the leading model in quantifying ecological resilience (Holling 1992), posits that organisms operate at specific spatial and temporal scales reflecting the availability of resources needed to support organisms’ survival and the ecological functions they perform. Quantifying redundancy and diversity of functions across these discontinuous scales, as well as the number of scales and cross-scale and within-scale diversity (Allen et al. 2005) allows for resilience estimate (Angeler et al. 2019, Roberts et al. 2019, Baho et al. 2020). Typically, redundancy and diversity of functions are derived from species traits (Craven et al. 2018, Gladstone-Gallagher et al. 2019), the phenotypic attributes of organisms that regulate their responses to the environment (*response traits*) and their effects on ecosystem processes (*effect traits*) (Truchy et al. 2015). Accordingly, redundancy among species having similar functional traits (i.e. functional redundancy) increases the buffering capacity of an ecosystem to stressors due to the higher likelihood of negatively affected species being replaced by functionally similar species (*the insurance hypothesis*, Naeem & Li 1997, Yachi & Loreau 1999). Response traits can be used to estimate the range of responses a functional group possesses against environmental change (i.e. response diversity, Elmqvist et al. 2003), meaning that if response diversity is low, many species will be similarly affected by a perturbation. Yet, studies testing the aforementioned metrics remain scarce (Jouffray et al. 2015, Angeler & Allen 2016), especially in dynamic socio-ecological systems such as rivers and streams (but see Pelletier et al. 2020).

Streams are often viewed as disturbance-prone ecosystems due to their high-variability in flow (Poff et al. 1997). Drying River Networks (DRN) experience periodic loss of surface water and dry in parts or along the whole stream channel (Datry et al. 2014a, Messager et al. 2021). By fragmenting the stream network (i.e. loss of connectivity between populations, communities and ecosystems, Cid et al. 2022), flow intermittence shapes community (i.e. community composition changes, loss of species richness, Datry et al. 2014a, Harris et al. 2018) and ecosystem functioning (Corti et al. 2011, Truchy et al. 2020) dynamics. Community responses depend on species resistance and resilience and the quality and accessibility of refuges (Sarremejane et al. 2021), reducing the long-term effects of flow intermittence on biodiversity and ecosystem functioning (Truchy et al. 2020, Sarremejane et al. 2024). For instance, drift (Vander Vorste et al. 2016a), vertical migration (Vander Vorste et al. 2016b), aerial colonization and resistance forms (Sarremejane et al. 2020) are all processes facilitating a quick return of a community to pre-disturbance composition (Vander Vorste et al. 2016c, Stubbington et al. 2024). Community composition can also be hardly altered by flow intermittence if they comprise stress-tolerant generalist taxa (Mykrä & Heino 2017). Yet, in other cases, resilience may be low if the access to refuges is severed, leading to negative impacts on ecosystem functioning (Datry et al. 2011, Truchy et al. 2020). Network position may also influence community responses to flow intermittence, with isolated headwater communities recovering more slowly than those in connected mainstem (Sarremejane et al. 2021).

In this study, we aim to explore how flow intermittence and stream network connectivity impact the resilience capacity of fish and invertebrate communities, and the subsequent consequences that this may have on ecosystem functioning. To do so, we monitored fish and invertebrate communities as well as a key function in stream ecosystems (leaf litter decay) in 20 sites across a river network naturally fragmented by drying. Using discontinuity analysis, indicators of resilience were inferred from the body size distribution of all individuals. Functional redundancy and response diversity were derived from species traits. Specifically, we tested the following hypotheses: (H1) as flow intermittence increases, the resilience capacity of communities lowers as non-perennial reaches exhibit less taxonomically and functionally diverse communities; (H2) resilience capacity may also change with stream network connectivity, with isolated upstream reaches having lower resilience than downstream reaches due to their naturally-reduced diversity and capacity to be recolonized. Finally, (H3) we expect a lower resilience capacity would translate into lower ecosystem functioning, i.e. lower decomposition rates, induced by a loss of functional redundancy or response diversity.

## Material and Methods

### Study sites

We monitored fish and instream invertebrate communities found in leaf litter at two sampling occasions (March-April and November-December 2021) to capture different life stages of invertebrates and fish, across 20 sites located in the Albarine river catchment (France; Datry 2012). The Albarine is subject to natural flow intermittence due to a permeable karstic bedrock and porous alluvial sediments that cause natural drying events lasting from weeks to months. The Albarine has a total network length of 150 kilometers of which >37% are known to be non-perennial from the headwaters to the main channel (Fig. 1). The 20 sampling sites were selected based on *a priori* knowledge of their flow regime (Datry et al. 2021) and represent a gradient of flow intermittence with an equal distribution of perennial and non-perennial reaches among upstream and downstream reaches (Fig. 1; Sarremejane et al. 2024, Silverthorne et al. 2024). At each non-perennial sampling site, we placed photo trap cameras set to take picture of the river channel twice-daily. From these pictures, we estimated the number of dry, non-flowing and flowing days and calculated flow intermittence, i.e. the proportion of non-flowing days during the study period (Mimeau et al. 2024, Truchy et al. 2023).

**Figure 1.**
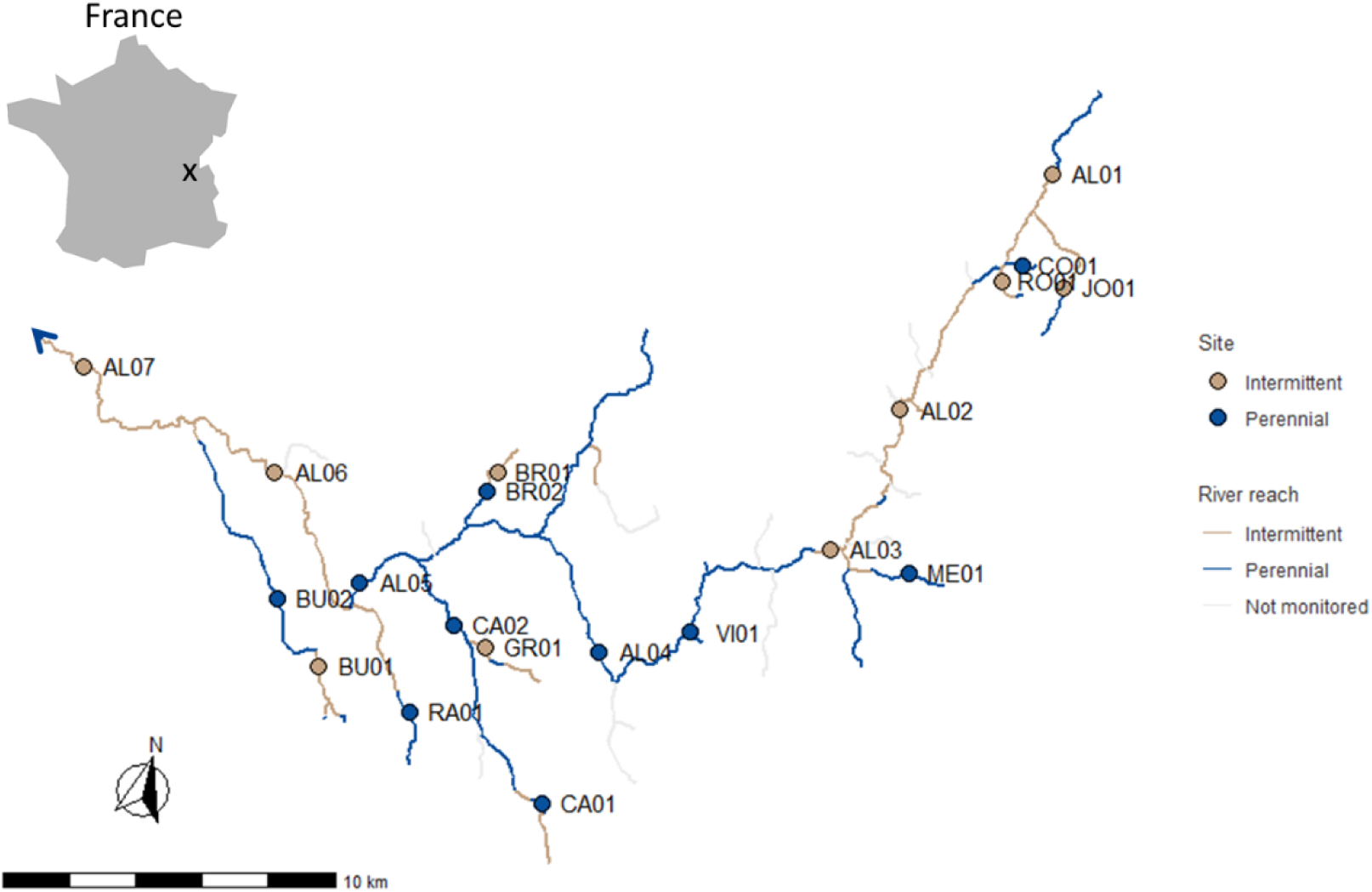
Map of the Albarine river catchment showing intermittent (orange) and perennial (blue) reaches (lines), sampling sites (dots) and their code names. Unmonitored reaches are indicated as grey lines. The blue arrow indicates flow direction.

### Environmental conditions

Using geographic information system (GIS) and digitalized maps of flow intermittence regimes across the Albarine catchment (Datry 2012, Gauthier et al. 2020), we calculated spatial variables representing site connectivity, including the distance to the source and the percentage of all perennial reaches upstream of each sampling site (i.e. upstream connectivity; see e.g. Sarremejane et al. 2024). Site altitude above sea level was also determined from a digitized elevation model.

At each sampling occasion and when water was present, we measured a set of environmental variables such as instream pH, conductivity and oxygen concentrations along with total dissolved nitrogen and phosphorus (HQ4300 with pH, conductivity and dissolved oxygen probes, Hach, USA), flow velocity (OTT MF pro, OTT HydroMet GmbH, Deutschland) and wetted width (averaged across 10 transects). Temperature was recorded hourly in the instream habitat using iButton loggers (Maxim Integrated Products, USA). Instream bryophyte and substrate (bedrock, boulders, cobbles, gravel and sand) covers were estimated visually. Instream sediment samples were dried (70⁰C) and burnt (550⁰C) to ashes in the laboratory to determine moisture and organic matter content, respectively. Instream canopy cover was assessed with a vertical tube (15 cm diameter) at 10 random locations. Environmental characteristics did not differ between non-perennial and perennial streams (Sarremejane et al. 2024) but differed between upstream and downstream reaches due to differences in mean wetted width, instream canopy cover, pH, soil moisture and organic matter contents (Sarremejane et al. 2024).

For graphical representation, sites were sometimes divided according to their network location (Fig. 1): headwaters (distance to source < 10km, n=14) and downstream reaches (distance to source > 10km, n=6).

### Leaf litter decomposition

Leaf litter decomposition was assessed by enclosing 4 ± 0.05 g (mean ± SD) of dried alder (*Alnus glutinosa*) leaves in 1 cm mesh bags, allowing for colonization by both microorganisms and invertebrates (Foulquier et al. 2015). Leaves were collected at abscission during autumn 2020, at a site approximately 80km away from the Albarine. For each field campaign, three litterbags were deployed in each stream for 29 ± 4 days. After retrieval, bags were taken to the laboratory and leaves were cleaned under tap water with invertebrates handpicked and stored in 70% ethanol for later identification. The remaining leaves were oven-dried for 78h at 70°C to estimate dry mass. A subsample was then ashed at 550°C to quantify ash free dry mass (AFDM). Breakdown rates were calculated for each litterbag based on the negative exponential decay model (K) corrected for the incubation time and cumulative temperatures (Benfield 1996). Decomposition rates were averaged across the six litterbags deployed at each site during year 2021. Of the 120 litterbags deployed, 6 were lost due to floods (Sarremejane et al. 2024).

### Invertebrate and fish communities

#### Sampling and identification

For each coarse-mesh litterbags, both aquatic and terrestrial invertebrates (size > 500 μm) were handpicked, identified under the microscope and counted in the lab (see Sarremejane et al. 2024). Taxa were identified at the finest taxonomic levels (Table S1), mostly genus and family. Terrestrial invertebrates were found colonizing litterbags during the dry phase in non-perennial reaches. Every individuals belonging to the same genus (or family, respectively) was photographed in a plate with graph paper underneath. Individual body size was estimated using the Segmented Line tool of ImageJ software version 1.53t (Rasband 1997).

Fish communities were sampled by two-pass removal electrofishing (CEN 2003). One to four fishing teams (each of 3 to 4 persons), depending on the stream width, waded in the upstream direction. Sampling was performed with upstream blocking nets, and covered areas ranging from 156 to 2780 m², depending on the site. All fish were identified, measured (total length in millimeters) and weighed (in grams) *in situ* and then released.

#### Functional traits

We assigned functional feeding groups (FFG) to each fish and invertebrate taxa according to their feeding habits (Tachet et al. 2010, Schmidt-Kloiber & Hering 2015). Feeding habits were divided as deposit-feeder, filter-feeder, grazer, insectivorous, parasite, piercer, piscivorous and shredder (Table S1). In case of variability in feeding habits (for invertebrates, in particular), taxa were assigned based on their dominant feeding habit. Information on feeding habits was missing for a total of 15 taxa representing 0.29% of the individuals and were thus not taken into account when calculating functional richness and redundancy (see below).

For both fish and invertebrates, we used a fuzzy-coding approach, with values of 1 to 3 indicating weak to strong affinity with each response trait category (i.e. pH preferences, trophic status, temperature preferences and current velocity preferences, Table S1 and references therein), that accounts for variability in preferences within taxa (e.g. current preferences may change during life cycles) and between species belonging to the same taxa. Response trait information was missing for 33% of the taxa – all of them being terrestrial invertebrate taxa – representing 320 individuals over the 13,846 collected individuals. We then transformed this information as percentage affinity for each response trait category and calculated, in each sample, the abundance of each category by multiplying each taxa percentage affinity by their abundance. pH preferences, trophic status, temperature preferences and current velocity preferences were chosen as they give a good representation of taxa habitat use.

### Cross-scale resilience metrics

Cross-scale resilience metrics are calculated by first identifying the discontinuous scales at which functions are performed by each species in the community (Nash et al. 2014) and then, using functional traits across scales to estimate functional redundancy and functional diversity within and across scales (Fischer et al. 2007).

#### Identifying discontinuous scales

We performed discontinuity analysis on invertebrate and fish community for each stream site and for both seasons combined to account for the overlapping life stages of a same species across time, using Bayesian Classification and Regression Trees (BCART; Stow et al. 2007). Body size is strongly allometric to the scale at which functions are performed by organisms (Holling 1992, Nash et al. 2014). Prior to the analysis, we prepared a univariate data matrix comprising of the ascending log-transformed body sizes. BCART (BCART software developed by Chipman et al. 1998) identifies groups in the body size data by assessing within-group homogeneity and is effective for identifying aggregations (Fig. S1; Bremner & Taplin 2004). The outcome is a branching tree, with the terminal nodes delineating aggregations of maximum homogeneity (Stow et al. 2007). Aggregations of maximum homogeneity comprise individuals similar in terms of body size that differ from other aggregations and presumably operate at distinct scales (Fig. S1).

From these analyses, for each sampling site, we obtained four site-specific metrics (Fig. S1):

- The number of scales, i.e. the number of body size aggregations;
- The within-scale richness estimated as the mean number of taxa present in each body size aggregation;
- The scale span which is the mean difference between the greatest and smallest log-transformed body size within the aggregation;
- The gap size evaluated as the mean distance in terms of log-transformed body size separating neighboring aggregations.

#### Estimating functional richness, cross- and within-scale functional redundancy

We determined functional richness by summing the number of functional feeding groups (FFG) present within each scale (i.e. body size aggregation) and averaging these numbers per site (Fig. S1). We also calculated within-scale functional redundancy as the number of taxa from each FFG within each body size aggregation (Peterson et al. 1998) and cross-scale functional redundancy as the number of aggregations for which each FFG has at least one representative (Fig. S1; Peterson et al. 1998).

Systems with high within-scale and cross-scale functional redundancy are the most resilient (Fig. S1; Allen et al. 2005, Biggs et al. 2020).

#### Estimating response diversity

For each FFG, we gathered the corresponding response traits, i.e. pH preferences, trophic status, temperature preferences and current velocity preferences (Table S1 and references therein). We quantified the response diversity of each FFG by measuring the multivariate functional dispersion (FDis; Laliberté & Legendre 2010) of the sampled taxa, using a Gower dissimilarity matrix. Since FDis is the average distance of individual species to the group centroid in response trait space, it is little influenced by species richness (Laliberté & Legendre 2010). Furthermore, the use of FDis in calculating response diversity ensures that response diversity measures are not trivially related to functional redundancy. If a site contained only one species of a particular FFG, no multivariate dispersion could be computed and a response diversity of 0 was assigned. Response diversity was not weighed by species abundances as rare species may contribute substantially to resilience (Walker et al. 1999). A decrease in response diversity for a given effect group means that its composition has shifted towards species that are more similar in how they respond to disturbance – indicating a loss of resilience.

### Statistical analyses

We designed linear models with resilience metrics, i.e. number of scales, within-scale richness, scale span, gap size, cross-scale and within-scale functional redundancy and, response diversity as response variables, respectively. Flow intermittence (H1), upstream distance to source and upstream proportion of perennial reaches (H2) and their interactions were fitted as explanatory variables. To test whether lower resilience capacity translates into lower ecosystem functioning (H3), we fitted linear models with leaf litter decomposition rates as response variable and cross-scale and within-scale functional redundancy as well as response diversity of shredders as explanatory variables, respectively. In few cases, response variables were log-transformed and outliers removed to ensure normal distribution of residuals and homoscedasticity. To reduce the disproportionate effect of a few large values of upstream distance to source, upstream distance to source was log-transformed in all models.

All linear models were implemented in R v 4.1.3 (R Core Team 2022) with the package vegan (Oksanen et al. 2022).

## Results

More than 13,800 individuals – belonging to 164 different taxa – were found and measured, of which 45.79% and 54.21% were fish and invertebrates, respectively. All functional feeding groups (FFG) were represented, with insectivorous and scraper fish being dominant in the fish community and, shredders and scrapers representing 71% of the invertebrate community.

### Effects of flow intermittence and network connectivity on cross-scale resilience metrics

Body sizes ranged from 0.062 to 275 mm in non-perennial reaches and from 0.055 to 538 mm in perennial reaches. Number of scales significantly decreased with flow intermittence (Fig. 2a; Table S2), with non-perennial reaches characterized by fewer scales than perennial reaches (non-perennial (NP): 9.78 ± 1.21; perennial reaches (P): 13.82 ± 1.48). This decrease in number of scales due to flow intermittence was stronger in upstream than downstream reaches (Fig. 2a; Table S2). Number of scales also increased with the % of upstream perennial reaches (upstream connectivity), but more strongly in downstream reaches (Fig. 2b; Table S2).

**Figure 2.**
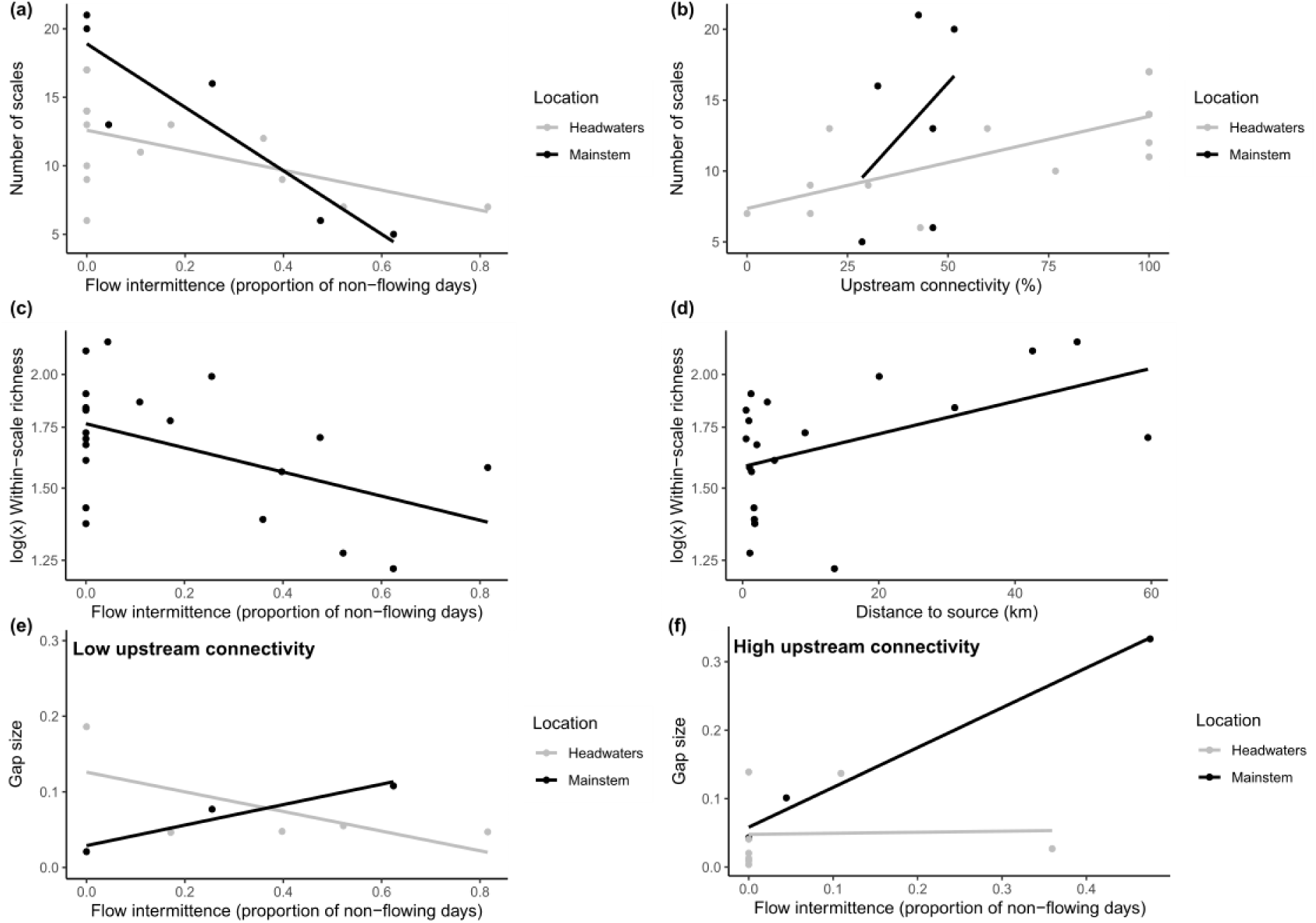
Responses of cross-scale resilience metrics – number of scales (a, b), within-scale richness (c, d), and gap size (e, f) to flow intermittence, upstream distance to source or upstream proportion of perennial reaches (upstream connectivity). Number of scales is the number of body size aggregations identified by the discontinuity analysis when applied to body size distribution. The within-scale richness is estimated as the number of taxa present in each body size aggregation. The distance in terms of body size separating two aggregations is called gap size. Responses in panels a, b, e and f were grouped by location (headwaters vs. mainsteam) to illustrate interactions. Panels e and f depict a three-way interactions: panel e presents sites with a lower upstream connectivity (i.e. upstream proportion of perennial reaches < 45%) while panel f presents sites with a higher upstream connectivity (i.e. upstream proportion of perennial reaches > 45%).

Within-scale richness decreased with flow intermittence (Fig. 2c; Table S2), whereas it increased with distance to source (Fig. 2d; Table S2). Non-perennial reaches had on average 8% fewer taxa than perennial reaches. Downstream reaches had on average 28% more taxa than upstream ones Scale span was only influenced by network connectivity, with shorter scales in highly connected upstream reaches and wider scales in highly connected downstream reaches (Table S2; Fig. S2).

Under low upstream connectivity conditions (UC < 45%), gap size in headwaters decreased with flow intermittence (Fig. 2e; Table S2). Gap size increased with flow intermittence in downstream reaches even more so under high upstream connectivity than under low upstream connectivity conditions (significant three-way interaction; F = 5.61; P = 0.04; Fig. 2e, f; Table S2).

### Effects of flow intermittence and network connectivity on functional richness, cross- and within-scale functional redundancy

Flow intermittence and network connectivity did not have any effects on functional richness. Patterns of cross-scale and within-scale functional redundancy for individual functional feeding groups included positive, negative and non-significant relationships with flow intermittence and upstream distance to source (Fig. 3). Non-perennial reaches had lower levels of cross-scale functional redundancy in filter-feeders, grazers, insectivores, piercers and shredders (Fig. 3a; Table S3).

**Figure 3.**
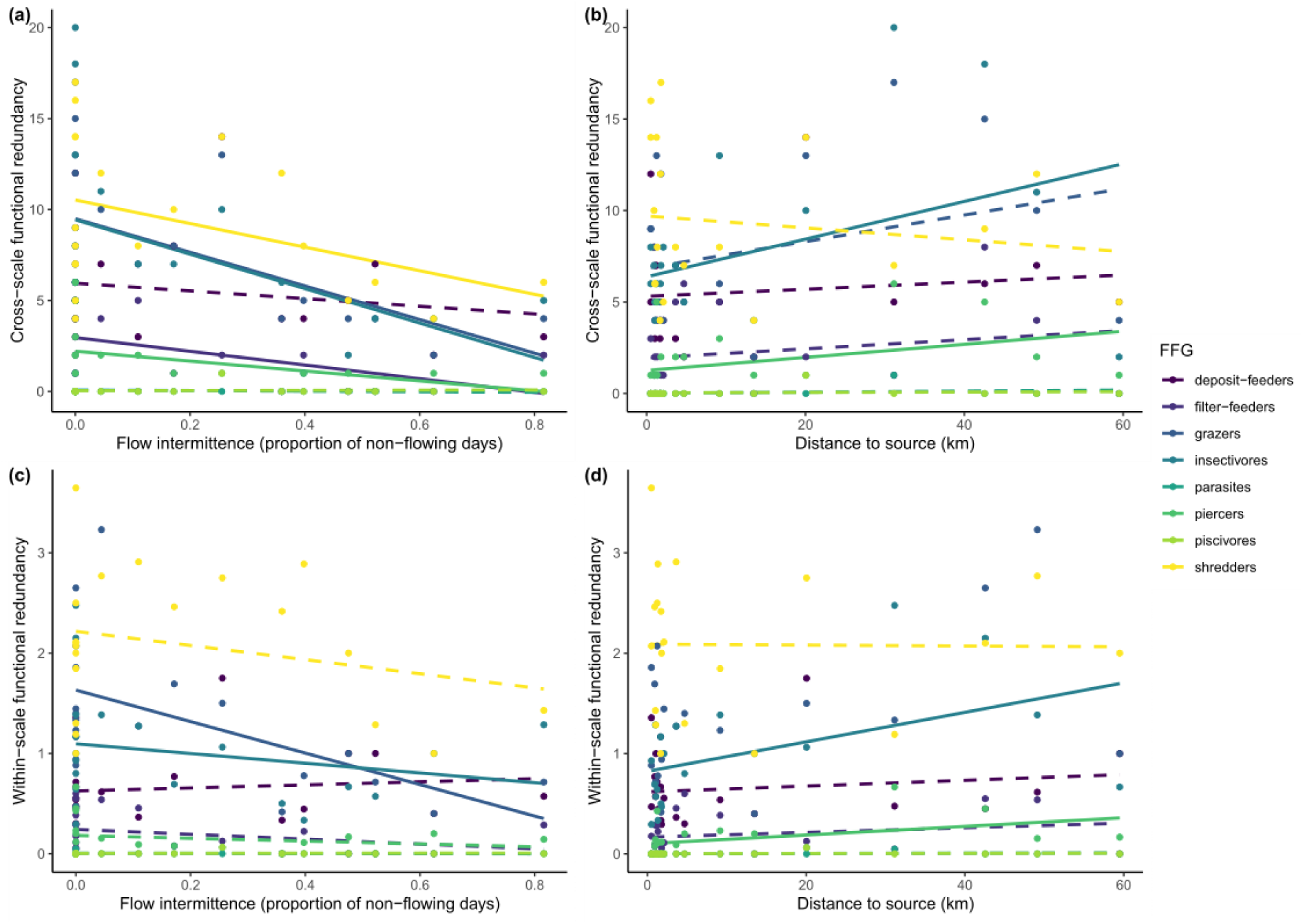
Responses of cross-scale (a, b) and within-scale (c, d) functional redundancy in eight functional feeding groups (FFG) to flow intermittence and upstream distance to source. Solid and dotted lines indicate significant and non-significant relationships, respectively. Statistics accompanying the models can be found in Tables S3-S4.

Headwaters had lower cross-scale functional redundancy in insectivores and piercers (Fig. 3b; Table S3). Cross-scale functional redundancy in shredders increased with upstream connectivity (Fig. S3; Table S3). Additionally, cross-scale functional redundancy in filter-feeders and insectivores decreased the most with flow intermittence in downstream reaches (significant two-way interactions: F > 6.92; P < 0.02; Fig. S4a, b; Table S3). Headwaters with low upstream connectivity conditions tended to have the lowest levels of cross-scale functional redundancy in grazers (Fig. S4c; Table S3).

Non-perennial reaches were characterized by lower levels of within-scale functional redundancy in grazers and insectivores (Fig. 3c; Table S4). Within-scale functional redundancy in insectivores and piercers was higher in downstream reaches (Fig. 3d; Table S4). Additionally, the highest levels of within-scale functional redundancy in filter-feeders and insectivores were observed in perennial downstream reaches (significant two-way interactions; Fig. S5; Table S4).

### Effects of flow intermittence and network connectivity on response diversity

Flow intermittence and network connectivity did not have any effects on the response diversity of the overall community. Non-perennial reaches were characterized by greater response diversity of shredders (Fig. 4a, Table S5). Response diversity of particular FFG increased with network connectivity, with greater response diversity of shredders in downstream reaches (Fig. 4b, Table S5) and greater response diversity of grazers at sites with higher proportions of upstream perennial reaches (Fig. 4c, Table S5). A three-way interaction indicated that under low upstream connectivity conditions (UC < 45%), response diversity of grazers decreased with flow intermittence, with the greatest decline in headwaters (Fig. S6; Table S5). Under high upstream connectivity, response diversity of grazers increased with flow intermittence, even more so in downstream reaches (F = 4.47; P = 0.05; Fig. S6; Table S5).

**Figure 4.**
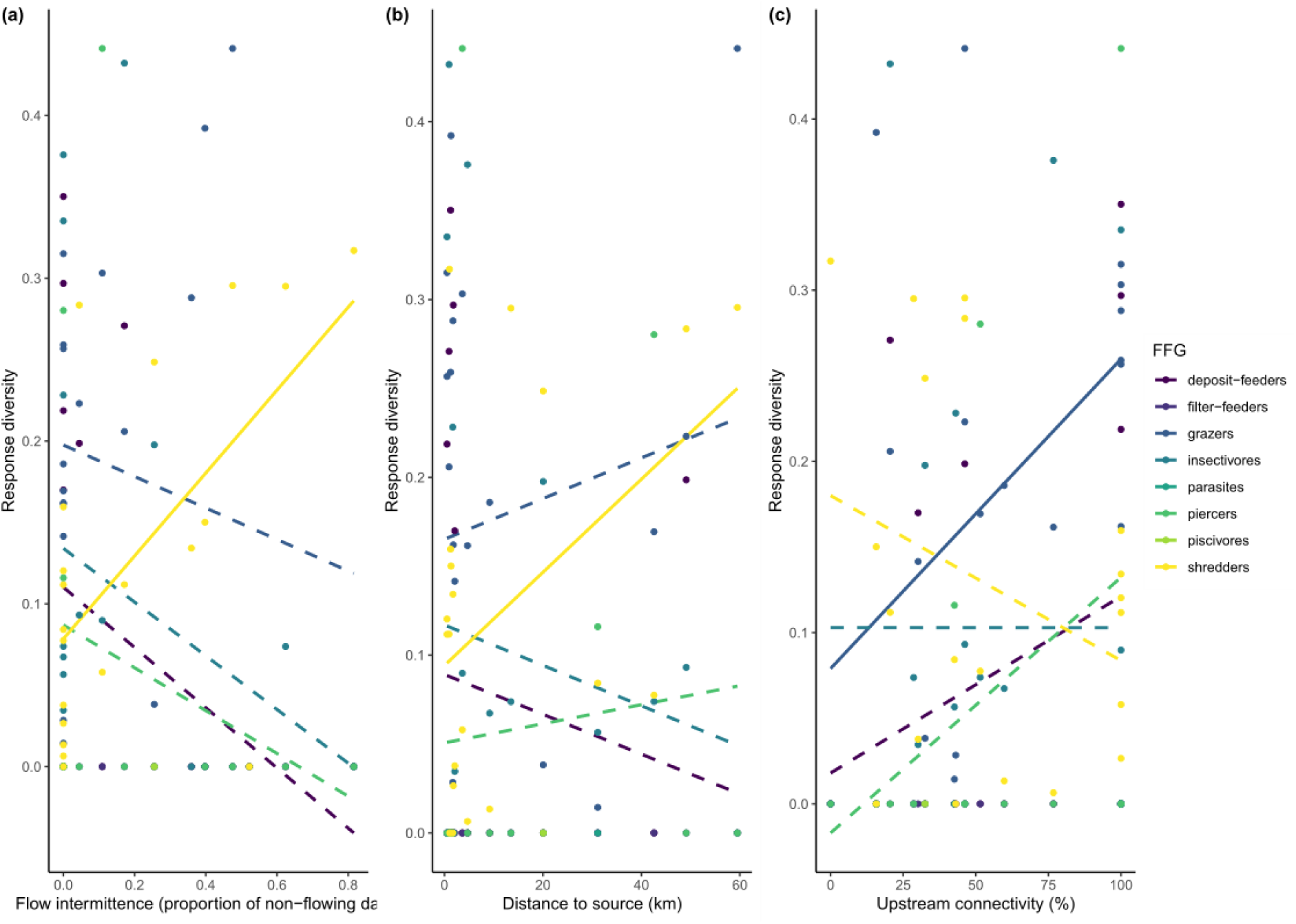
Responses diversity (a-c) of eight functional feeding groups (FFG) to flow intermittence (a), upstream distance to source (b) and upstream proportion of perennial reaches (upstream connectivity, c). Relationships for some FFG were not testable as response diversity was not computable in some of the sites (see Methods for details). Solid and dotted lines indicate significant and non-significant relationships, respectively. Statistics accompanying the models can be found in Table S5.

### Resilience capacity and ecosystem functioning (H4)

In perennial reaches, litter decomposition rates decreased with greater cross-scale and within-scale functional redundancy in shredders (F = 7.93; P = 0.02 and F = 9.70; P = 0.01, respectively; Fig. 5a-b), as well as with greater response diversity of shredders (F = 10.85; P = 0.009; Fig 5c). None of these relationships were significant in non–perennial reaches (Fig. 5).

**Figure 5.**
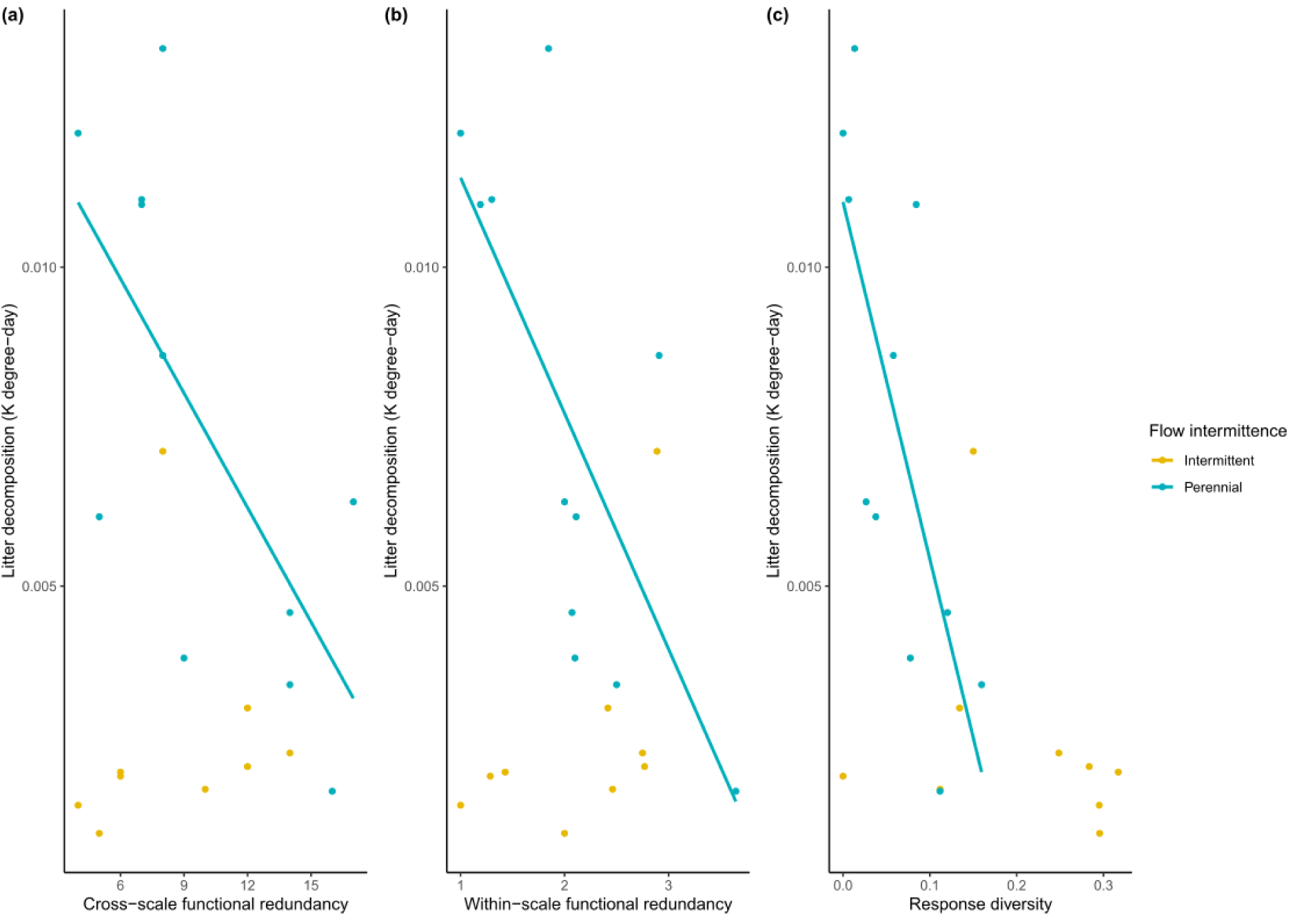
Relationships between yearly-averaged litter decomposition rates in coarse-mesh bags, from both non-perennial (yellow) and perennial (blue) reaches, and resilience metrics, i.e. cross-scale functional redundancy (a), within-scale functional redundancy (b) and response diversity (c) of shredder communities. Solid lines indicate significant relationships.

## Discussion

Numerous studies have characterized the ecological responses of communities to drying and connectivity patterns in river networks, showing greater compositional shifts and decreases in taxonomic richness, recovery and abundance in response to drying when connectivity is low (Bogan et al. 2013, Datry et al. 2014b, Jacquet et al. 2022, Datry et al. 2023). However, assessments of ecological resilience (*sensu* Holling, 1973) are missing from drying river networks. Our study explored community resilience capacity in fragmented river networks, which is becoming critical as drying intensifies with global change (Zipper et al. 2021, Datry et al. 2023). Our results indicate that flow intermittence decreases the resilience capacity of both fish and invertebrate communities, as predicted. (H1). We also found that upstream reaches have lower resilience capacity than downstream reaches. Finally, and contrary to our expectations, the loss of resilience capacity did not necessarily translates into ecosystem functioning alteration due to the high functional redundancy across fish and invertebrate communities.

Flow intermittence, alone, led to a decrease in four resilience indicators out of seven. Non-perennial reaches were characterized by lower number of scales that each contain fewer species, in accordance with previous studies showing a lower taxonomic richness (Datry et al. 2014a, Crabot et al. 2021, Yang et al. 2023, Gianuca et al. 2024). Selection of a subset of species with drying resistance strategies may limit functional redundancy (i.e. dominance of drying-resistant species, Hill et al. 2019, Bozóki et al. 2024). Fish population, on the contrary, are especially sensitive to drying (Davey & Kelly 2007, Benejam et al. 2010), leading to a complete absence of fish in strongly non-perennial reaches. During drying events, fish can survive in deep pools as long as oxygen conditions remain satisfying (Hodges & Magoulick 2011, Gilmore et al. 2018, Hill & Milner 2018), allowing them to recolonize when flow resumes (Pires et al. 2014, Driver & Hoeinghaus 2016). However, the presence of pools is very limited in space and time in the Albarine (Sarremejane et al. 2024, Silverthorn et al. 2024). The important community shift translates to an overall decrease in functional redundancy – a result in contradiction with previous studies (Crabot et al. 2021, Chanut et al. 2023, Gianuca et al. 2024, Escobar-Camacho et al. 2025). Our study shows a significant decrease in cross-scale functional redundancy of five functional feeding groups out of the eight in response to increasing flow intermittency. Cross-scale functional redundancy of filter-feeders, grazers, insectivores, piercers and shredders were negatively affected by flow intermittence. Together, these results mean that non-perennial reaches are characterized by fewer functionally-similar species/individuals, with some scales completely lacking representatives of some functional groups.

The insurance hypothesis (Naeem & Li 1997, Yachi & Loreau 1999), predicts that a loss of functional redundancy reduces the buffering capacity of an ecosystem to stressors, potentially impairing ecosystem functioning. However, we did not find any negative relationship between (cross-scale or within-scale) functional redundancy of shredders and the measured rates of leaf litter decomposition in non-perennial reaches. In these reaches, despite high levels of response diversity of shredders, leaf litter decomposition rates were particularly low, potentially reflecting shredder communities that are still recovering from drying and are therefore not fully operational yet for processing such an abundant resource (Sarremejane et al. 2024). This apparent lack of litter processing in non-perennial reaches may have further impacts on river carbon budget (Datry et al. 2018). However, litter decomposition rates slowed down with increasing cross-scale and within-scale functional redundancy of shredders. Such results may indicate 1) the dominance of a very competitive and efficient decomposer at some perennial sites and 2) a competition between the many co-occurring shredder taxa that tend to slow down leaf litter decomposition rates. Low quantity of leaves in these systems combined with high invertebrate densities (Sarremejane et al. 2024) may promote aggregation of shredders on leaf packs, leading to a higher interspecific competition that reduces shredder efficiency at decomposing leaf litter (Cardinale, 2011, Firmino et al. 2022). The perennial reaches of the Albarine catchment exhibiting the highest leaf litter decomposition rates were dominated by *Gammarus* sp., a very effective decomposer (Dangles & Malmqvist, 2004) and a strong competitor that might outcompete other shredder species, hence the low functionalredundancy and response diversity observed in these sites.

As multiple small streams form larger rivers, the dendritic structure of river networks can promote increases in local metrics of biodiversity (e.g. Finn et al. 2011, Göthe et al. 2014a, Jyrkänkallio-Mikkola et al. 2018) and productivity (Ferreira et al. 2023) as influx of organisms and resources (e.g. nutrients and dissolved organic matter) increase from up to downstream (Vannote et al. 1980). In contrast, headwater streams are generally prone to a very high regional beta-diversity. This longitudinal connectivity is one of the fundamental dimensions within watersheds (Townsend 1989, Ward 1989) and imply downstream ecosystems rely on the connectivity with upstream ecosystems. Accordingly, headwaters were characterized by lower levels of within-scale richness, lower levels of cross-scale and within-scale functional redundancy in insectivores and piercers as well as lower response diversity of shredders, which is in accordance with previous studies showing overall low redundancy in headwaters (Göthe et al. 2014b). These results are not unexpected as headwater streams are usually characterized by low alpha diversity (but not necessarily low beta diversity, Richardson 2019), resulting largely from environmental heterogeneity and their isolated position in the river network. Isolation and strong environmental filtering prevents some species from colonizing and establishing (dispersal limitation, Leibold et al. 2004, Sarremejane et al. 2017). Surprisingly, flow intermittence affected more strongly invertebrate community resilience in down-than in upstream reaches. This observed pattern could reflect 1) the effects of an upstream perturbation regime that accumulate along the river network and 2) strong contrasted diversity patterns between perennial and non-perennial reaches. Forested buffer strips along rivers play an important role in shading, cooling and keeping moisture in the aquatic environment (Cole et al. 2020, Sargac et al. 2021). These positive effects are particularly important in non-perennial headwaters (i.e. narrower river bed of upstream reaches compared to downstream reaches) and could promote the survival of more taxa. After a drying event, community recovery is indeed expected to be slower in headwaters in comparison with downstream sites, where recolonization rates are considerably higher due to mass effect (Leibold et al. 2004, Brown et al. 2011). Dispersal traits i.e. the ability of species to recolonize dry reaches from perennial ones after flow resumes (Gauthier et al. 2020, Sarremejane et al. 2020) and dispersal distances that can be covered by individuals within the river network are also parameters that should be considered in the context of recolonization after a disturbance. Species with high fecundity, small body size, short life span, high dispersal ability and dormancy, should be selected in non-perennial river networks (Vander Vorste et al. 2016a, 2016b, Sarremejane et al. 2020). Unfortunately, we could not include these traits in our assessment of response diversity as trait values were available for aquatic invertebrates (Sarremejane et al. 2020) but not for fish. It could have been informative to assess whether dispersal-related traits affected community response diversity in relation with drying intensity and network fragmentation.

The effects of flow intermittence on stream communities and functioning are relatively well documented (Datry et al. 2011, Soria et al. 2017, Crabot et al. 2021, Chanut et al. 2023, Escobar-Camacho et al. 2025) whereas the effects of network fragmentation remains overlooked. Although some evidence exists for fish (Jaeger et al. 2014), invertebrate community richness (Gauthier et al. 2020, Crabot et al. 2021, Sarremejane et al. 2021) and leaf-litter decomposition (Abril et al. 2016, Sarremejane et al. 2024), little is known about the effects of fragmentation on community resilience capacity. The proportion of perennial reaches upstream increased cross-scale functional redundancy in shredders and response diversity in grazers. In order for an ecosystem function provided by a certain functional group to be resilient to an, array of disturbances, there must be high levels of both functional redundancy and response diversity of that group (Nyström 2006). Along the network fragmentation gradient, the decoupling between functional redundancy and response diversity we observed for both shredders and grazers suggests that the functions these groups perform may still have some level of resilience due to redundancy in shredders and differences in response diversity of grazers, respectively. Nevertheless, network fragmentation may still threaten both grazer and shredder communities, and the functions they provide – specifically the transfer of carbon through the food web (Ledger et al. 2013, Lu et al. 2016) – when facing other pressures, such as water pollution. Greater amount of network fragmentation (i.e. proportion of perennial reaches upstream < 45%) indeed resulted in lower response diversity of grazer communities when exposed to flow intermittence. Perennial reaches upstream of a non-perennial reach may promote 1) growth pulses of high-quality biofilm and 2) the arrival of diverse adapted grazer species, after rewetting events (Timoner et al. 2012, 2014).

Resilience is an emergent property of complex systems and no single metric can perfectly describe it (Angeler & Allen 2016). The assessment of scaling patterns in ecosystems is important because organisms operate at specific spatial and temporal scales that reflect the availability of resources needed for organism’ survival and the ecological functions they perform. Resilience can therefore be assessed by 1) considering simultaneously multiples metrics (i.e. number, within-scale richness, span, gap size – the *cross-scale resilience model*, Allen et al. 2005, Angeler & Allen 2016, Roberts et al. 2019) and 2) combining it with functional redundancy and response diversity within and across scales. We show that the responses of individual resilience metrics varied with flow intermittence and network connectivity. Flow intermittence altered the overall resilience capacity of invertebrate and fish communities by fragmenting the river network, with negative impacts on ecosystem functioning being apparent. Headwater and non-perennial reaches appear to be particularly at risk due to their isolation and existing environmental filters that prevent recolonization. In order to progress resilience theory, comparing standardized resilience metrics between systems is required. It remains unclear – for stream ecosystems – under which conditions resilience capacity is eroded to such an extent that regime shifts would occur.

## Acknowledgements

We thank Angélique Arbaretaz, Abdel Azougui, Maxence Forcellini, Margot Jans, Bertrand Launay and Guillaume Le Goff for their help in the lab and in the field. We also thank Etienne Dambrine for collecting the leaf litter and the land owners (including Didier Cristini) on the Albarine catchment, who provided access to their properties. This research was supported by the European Union’s Horizon 2020 research and innovation programme under the Marie Skłodowska-Curie grant agreement no. 891090 (MetaDryNet) and the DRYvER project (Datry et al. 2021; grant agreement no. 869226).

## Notes

### Competing Interest Statement

The authors have declared no competing interest.

