## Supplementary file for "Community resilience in a river network: the roles of connectivity and drying regime"

**Table S1.** Taxa list from instream habitats, including invertebrate and fish taxa. Their total abundance and their trait information – feeding habits – are also reported.

| Typical habitat | Order | Family | Lowest taxonomic level | Total abundance | absorber | deposit feeder/humus feeder | shredder / decomposer | scraper / grazer | filter-feeder | piercer | insectivorous | piscivorous | parasite | Reference used for traits |
|---|---|---|---|---|---|---|---|---|---|---|---|---|---|---|
| Aquatic | Cypriniformes | Nemacheilidae | Barbatula barbatula | 890 | 0 | 0 | 0 | 0 | 0 | 0 | 1 | 1 | 0 | 1 |
| Aquatic | Cypriniformes | Cyprinidae | Blicca bjoerkna | 3 | 0 | 0 | 0 | 0 | 0 | 0 | 0 | 0 | 0 | 1 |
| Aquatic | Scorpaeniformes | Cottidae | Cottus gobio | 571 | 0 | 0 | 0 | 0 | 0 | 0 | 1 | 0 | 0 | 1 |
| Aquatic | Perciformes | Centrarchidae | Lepomis gibbosus | 11 | 0 | 0 | 0 | 0 | 0 | 0 | 1 | 1 | 0 | 1 |
| Aquatic | Cypriniformes | Cyprinidae | Squalius cephalus | 6 | 0 | 0 | 0 | 1 | 0 | 0 | 1 | 1 | 0 | 1 |
| Aquatic | Cypriniformes | Cyprinidae | Telestes souffia | 11 | 0 | 0 | 0 | 1 | 0 | 0 | 1 | 0 | 0 | 1 |
| Aquatic | Cypriniformes | Cyprinidae | Phoxinus phoxinus | 750 | 0 | 0 | 1 | 1 | 0 | 0 | 1 | 0 | 0 | 1 |
| Aquatic | Cypriniformes | Cyprinidae | Pseudorasbora parva | 3 | 0 | 0 | 0 | 0 | 0 | 0 | 0 | 0 | 0 | 1 |
| Aquatic | Cypriniformes | Cyprinidae | Rutilus rutilus | 6 | 0 | 0 | 0 | 0 | 0 | 0 | 1 | 0 | 0 | 1 |
| Aquatic | Salmoniformes | Salmonidae | Salmo trutta | 588 | 0 | 0 | 0 | 0 | 0 | 0 | 1 | 1 | 0 | 1 |
| Aquatic | Cypriniformes | Cyprinidae | Scardinius erythrophthalmus | 3 | 0 | 0 | 0 | 1 | 1 | 0 | 1 | 0 | 0 | 1 |
| Aquatic | Salmoniformes | Salmonidae | Thymallus thymallus | 167 | 0 | 0 | 0 | 0 | 0 | 0 | 1 | 0 | 0 | 1 |
| Aquatic | Cypriniformes | Cyprinidae | Tinca tinca | 2 | 0 | 0 | 1 | 1 | 0 | 0 | 1 | 0 | 0 | 1 |
| Aquatic | Acarina | | Hydracarina | 24 | 0 | 0 | 0 | 0 | 0 | 3 | 0 | 0 | 1 | 2 |
| Aquatic | Amphipoda | Gammaridae | Gammarus sp. | 3063 | 0 | 0 | 3 | 1 | 0 | 0 | 0 | 0 | 0 | 3 |
| Aquatic | Achaeta | Glossiphoniidae | Erpobdella sp. | 5 | 0 | 0 | 0 | 0 | 0 | 0 | 3 | 0 | 0 | 3 |
| Aquatic | Achaeta | Glossiphoniidae | Glossiphonia sp. | 2 | 0 | 0 | 0 | 0 | 0 | 3 | 0 | 0 | 1 | 3 |
| Aquatic | Achaeta | Glossiphoniidae | Helobdella stagnalis | 1 | 0 | 0 | 0 | 0 | 0 | 3 | 0 | 0 | 0 | 3 |
| Aquatic | Amphipoda | Niphargidae | Niphargus sp. | 3 | 0 | 0 | 3 | 0 | 0 | 0 | 0 | 0 | 0 | 3 |
| Aquatic | Achaeta | Piscicolidae | Piscicola sp. | 7 | 0 | 0 | 0 | 0 | 0 | 2 | 0 | 0 | 2 | 3 |
| Aquatic | Coleoptera | Chrysomelidae | Chrysomelidae | 8 | 0 | 0 | 0 | 3 | 0 | 0 | 0 | 0 | 0 | 3 |
| Aquatic | Coleoptera | Dryopidae | Pomatinus substriatus | 6 | 0 | 0 | 3 | 3 | 0 | 0 | 0 | 0 | 0 | 3 |
| Aquatic | Coleoptera | Dytiscidae | Agabus sp. | 11 | 0 | 0 | 3 | 0 | 0 | 3 | 0 | 0 | 0 | 3 |
| Aquatic | Coleoptera | Dytiscidae | Coelambus sp. | 1 | NA | NA | NA | NA | NA | NA | NA | NA | NA | NA |
| Aquatic | Coleoptera | Dytiscidae | Oreodytes sp. | 1 | 0 | 0 | 3 | 0 | 0 | 3 | 0 | 0 | 0 | 3 |
| Aquatic | Coleoptera | Elmidae | Elmis sp. | 4 | 0 | 0 | 2 | 3 | 0 | 0 | 0 | 0 | 0 | 3 |
| Aquatic | Coleoptera | Elmidae | Esolus sp. | 2 | 0 | 0 | 1 | 3 | 0 | 0 | 0 | 0 | 0 | 3 |
| Aquatic | Coleoptera | Elmidae | Limnius sp. | 3 | 0 | 0 | 2 | 3 | 0 | 0 | 0 | 0 | 0 | 3 |
| Aquatic | Coleoptera | Elmidae | Riolus sp. | 1 | 0 | 0 | 1 | 3 | 0 | 0 | 0 | 0 | 0 | 3 |
| Aquatic | Coleoptera | Haliplidae | Haliplus sp. | 1 | 0 | 0 | 1 | 3 | 0 | 3 | 0 | 0 | 0 | 3 |
| Aquatic | Coleoptera | Helophoridae | Helophorus sp. | 2 | 0 | 0 | 1 | 3 | 0 | 0 | 0 | 0 | 0 | 3 |
| Aquatic | Coleoptera | Hydraenidae | Hydraena sp. | 63 | 0 | 0 | 1 | 3 | 0 | 0 | 0 | 0 | 0 | 3 |
| Aquatic | Coleoptera | Hydrophilidae | Coelostoma sp. | 5 | 0 | 0 | 3 | 3 | 0 | 0 | 0 | 0 | 0 | 3 |
| Aquatic | Coleoptera | Hydroporinae | Deronectes sp. | 2 | 0 | 0 | 3 | 0 | 0 | 3 | 0 | 0 | 0 | 3 |
| Aquatic | Coleoptera | Hygrobiidae | Hygrobia hermanni | 1 | 0 | 0 | 1 | 0 | 0 | 0 | 3 | 0 | 0 | 3 |
| Aquatic | Coleoptera | Scirtidae | Elodes sp. | 50 | 0 | 0 | 1 | 3 | 0 | 0 | 0 | 0 | 0 | 3 |
| Aquatic | Coleoptera | Scirtidae | Odeles sp. | 16 | 0 | 0 | 1 | 3 | 0 | 0 | 0 | 0 | 0 | 3 |
| Aquatic | Diptera | Athericidae | Atherix sp. | 31 | 0 | 0 | 0 | 0 | 0 | 3 | 0 | 0 | 0 | 3 |

| | | | | | | | | | | | | | | |
|---|---|---|---|---|---|---|---|---|---|---|---|---|---|---|
| Aquatic | Diptera | Ceratopogonidae | Ceratopogoninae | 1 | 0 | 2 | 1 | 1 | 0 | 0 | 1 | 0 | 0 | 3 |
| Aquatic | Diptera | Chironomidae | Chironomini | 86 | 0 | 3 | 2 | 1 | 2 | 0 | 1 | 0 | 1 | 3 |
| Aquatic | Diptera | Chironomidae | Corinoneura sp. | 4 | 0 | 1 | 0 | 3 | 1 | 0 | 0 | 0 | 1 | 3 |
| Aquatic | Diptera | Chironomidae | Diamesinae | 14 | 0 | 1 | 0 | 3 | 1 | 0 | 0 | 0 | 1 | 3 |
| Aquatic | Diptera | Chironomidae | Orthocladiinae | 933 | 0 | 1 | 0 | 3 | 1 | 0 | 0 | 0 | 1 | 3 |
| Aquatic | Diptera | Chironomidae | Prodiamesinae | 1 | 0 | 1 | 0 | 3 | 1 | 0 | 0 | 0 | 1 | 3 |
| Aquatic | Diptera | Chironomidae | Tanypodinae | 52 | 0 | 0 | 0 | 0 | 0 | 0 | 3 | 0 | 0 | 3 |
| Aquatic | Diptera | Chironomidae | Tanytarsini | 62 | 0 | 3 | 2 | 2 | 1 | 0 | 0 | 0 | 0 | 3 |
| Aquatic | Diptera | Dixidae | Dixa sp. | 10 | 0 | 0 | 1 | 0 | 3 | 0 | 1 | 0 | 0 | 3 |
| Aquatic | Diptera | Empididae | Empididae | 3 | 0 | 0 | 0 | 0 | 0 | 0 | 3 | 0 | 0 | 3 |
| Aquatic | Diptera | Limoniidae | Limoniidae | 22 | 0 | 2 | 2 | 1 | 0 | 0 | 2 | 0 | 0 | 3 |
| Aquatic | Diptera | Psychodidae | Psychodidae | 4 | 0 | 2 | 3 | 1 | 0 | 0 | 0 | 0 | 0 | 3 |
| Aquatic | Diptera | Rhagionidae | Rhagionidae | 1 | 0 | 0 | 0 | 0 | 0 | 3 | 0 | 0 | 0 | 3 |
| Aquatic | Diptera | Sciomyzidae | Sciomyzidae | 2 | 0 | 0 | 0 | 0 | 0 | 3 | 0 | 0 | 2 | 3 |
| Aquatic | Diptera | Simuliidae | Simuliidae | 243 | 0 | 0 | 0 | 1 | 3 | 0 | 0 | 0 | 0 | 3 |
| Aquatic | Diptera | Stratiomyidae | Oxycera sp. | 5 | 0 | 2 | 3 | 1 | 0 | 0 | 1 | 0 | 0 | 3 |
| Aquatic | Diptera | Tipulidae | Tipulidae | 5 | 0 | 2 | 3 | 0 | 0 | 0 | 2 | 0 | 0 | 3 |
| Aquatic | Ephemeroptera | Baetidae | Alainites sp. | 58 | 0 | 1 | 0 | 3 | 0 | 0 | 0 | 0 | 0 | 3 |
| Aquatic | Ephemeroptera | Baetidae | Baetis sp. | 47 | 0 | 1 | 0 | 3 | 0 | 0 | 0 | 0 | 0 | 3 |
| Aquatic | Ephemeroptera | Baetidae | Centroptilum luteolum | 1 | 0 | 2 | 0 | 3 | 0 | 0 | 0 | 0 | 0 | 3 |
| Aquatic | Ephemeroptera | Ephemerellidae | Serratella ignita | 110 | 0 | 1 | 2 | 2 | 0 | 0 | 1 | 0 | 0 | 3 |
| Aquatic | Ephemeroptera | Ephemerellidae | Torleya major | 5 | 0 | 0 | 0 | 3 | 0 | 0 | 0 | 0 | 0 | 3 |
| Aquatic | Ephemeroptera | Ephemeridae | Ephemera sp. | 3 | 0 | 1 | 3 | 0 | 3 | 0 | 1 | 0 | 0 | 3 |
| Aquatic | Ephemeroptera | Heptageniidae | Ecdyonurus sp. | 60 | 0 | 0 | 2 | 3 | 0 | 0 | 0 | 0 | 0 | 3 |
| Aquatic | Ephemeroptera | Heptageniidae | Electrogena sp. | 2 | 0 | 0 | 0 | 3 | 0 | 0 | 0 | 0 | 0 | 3 |
| Aquatic | Ephemeroptera | Heptageniidae | Epeorus sp. | 3 | 0 | 1 | 0 | 3 | 0 | 0 | 0 | 0 | 0 | 3 |
| Aquatic | Ephemeroptera | Heptageniidae | Rhithrogena sp. | 4 | 0 | 1 | 0 | 3 | 0 | 0 | 0 | 0 | 0 | 3 |
| Aquatic | Ephemeroptera | Leptophlebiidae | Habroleptoides sp. | 431 | 0 | 0 | 2 | 3 | 0 | 0 | 0 | 0 | 0 | 3 |
| Aquatic | Ephemeroptera | Leptophlebiidae | Habrophlebia sp. | 40 | 0 | 1 | 3 | 0 | 0 | 0 | 0 | 0 | 0 | 3 |
| Aquatic | Ephemeroptera | Leptophlebiidae | Paraleptophlebia sp. | 3 | 0 | 2 | 2 | 1 | 0 | 0 | 0 | 0 | 0 | 3 |
| Aquatic | Gasteropoda | Lymnaeidae | Galba truncatula | 2 | 0 | 0 | 1 | 3 | 0 | 0 | 0 | 0 | 0 | 3 |
| Aquatic | Gasteropoda | Lymnaeidae | Omphiscola sp. | 1 | 0 | 0 | 1 | 3 | 0 | 0 | 0 | 0 | 0 | 3 |
| Aquatic | Gasteropoda | Lymnaeidae | Physa sp. | 5 | 0 | 0 | 1 | 3 | 0 | 0 | 0 | 0 | 0 | 3 |
| Aquatic | Gasteropoda | Lymnaeidae | Radix sp. | 4 | 0 | 0 | 1 | 3 | 0 | 0 | 0 | 0 | 0 | 3 |
| Aquatic | Gasteropoda | Planorbidae | Gyraulus sp. | 2 | 0 | 0 | 1 | 3 | 0 | 0 | 0 | 0 | 0 | 3 |
| Aquatic | Hemiptera | Veliidae | Mesovelia sp. | 1 | 0 | 0 | 0 | 0 | 0 | 3 | 0 | 0 | 0 | 3 |
| Aquatic | Hemiptera | Veliidae | Velia sp. | 1 | 0 | 0 | 0 | 0 | 0 | 3 | 1 | 0 | 0 | 3 |
| Aquatic | Hymenoptera | Agriotypidae | Agriotypus armatus | 3 | 0 | 0 | 0 | 0 | 0 | 0 | 0 | 0 | 3 | 3 |
| Aquatic | Isopoda | Asellidae | Asellus aquaticus | 17 | 0 | 0 | 3 | 0 | 0 | 0 | 0 | 0 | 0 | 3 |
| Aquatic | Plecoptera | Capniidae | Capnioneura sp. | 3 | 0 | 0 | 3 | 0 | 0 | 0 | 0 | 0 | 0 | 3 |
| Aquatic | Plecoptera | Leuctridae | Leuctra sp. | 222 | 0 | 1 | 3 | 1 | 0 | 0 | 0 | 0 | 0 | 3 |
| Aquatic | Plecoptera | Nemouridae | Amphinemura sp. | 128 | 0 | 1 | 3 | 0 | 0 | 0 | 0 | 0 | 0 | 3 |
| Aquatic | Plecoptera | Perlidae | Perla marginata | 2 | 0 | 0 | 3 | 0 | 0 | 0 | 1 | 0 | 0 | 3 |
| Aquatic | Plecoptera | Taeniopterygidae | Brachyptera sp. | 7 | 0 | 0 | 1 | 3 | 0 | 0 | 0 | 0 | 0 | 3 |
| Aquatic | Plecoptera | Perlodidae | Isoperla sp. | 14 | 0 | 0 | 3 | 1 | 0 | 0 | 1 | 0 | 0 | 3 |
| Aquatic | Plecoptera | Nemouridae mouridae | Nemoura sp. | 303 | 0 | 0 | 3 | 0 | 0 | 0 | 0 | 0 | 0 | 3 |
| Aquatic | Plecoptera | Perlodidae | Perlodes sp. | 32 | 0 | 0 | 3 | 0 | 0 | 0 | 1 | 0 | 0 | 3 |

| | | | | | | | | | | | | | | |
|---|---|---|---|---|---|---|---|---|---|---|---|---|---|---|
| Aquatic | Plecoptera | Nemouridae | Protonemura sp. | 9 | 0 | 0 | 3 | 0 | 0 | 0 | 0 | 0 | 0 | 3 |
| Aquatic | Plecoptera | Taeniopterygidae | Taeniopteryx schoenemundi | 1 | 0 | 0 | 3 | 0 | 0 | 0 | 0 | 0 | 0 | 3 |
| Aquatic | Plecoptera | Capniidae | Zwicknia sp. | 6 | 0 | 0 | 3 | 0 | 0 | 0 | 0 | 0 | 0 | 3 |
| Aquatic | Trichoptera | Glossosomatidae | Synagapetus sp. | 1 | 0 | 0 | 0 | 3 | 0 | 0 | 0 | 0 | 0 | 3 |
| Aquatic | Trichoptera | Hydropsychidae | Hydropsyche sp. | 35 | 0 | 0 | 0 | 0 | 3 | 0 | 1 | 0 | 0 | 3 |
| Aquatic | Trichoptera | Lepidostomatidae | Lepidostoma hirtum | 6 | 0 | 0 | 3 | 1 | 0 | 0 | 0 | 0 | 0 | 3 |
| Aquatic | Trichoptera | Limnephilidae | Allogamus/Melampop hylax | 10 | 0 | 0 | 3 | 0 | 0 | 0 | 0 | 0 | 0 | 3 |
| Aquatic | Trichoptera | Limnephilidae | Anabolia sp. | 1 | 0 | 1 | 3 | 2 | 0 | 0 | 1 | 0 | 0 | 3 |
| Aquatic | Trichoptera | Limnephilidae | Drusus sp. | 1 | 0 | 0 | 2 | 3 | 0 | 0 | 0 | 0 | 0 | 3 |
| Aquatic | Trichoptera | Limnephilidae | Glyphotaelius sp. | 5 | 0 | 0 | 3 | 0 | 0 | 0 | 0 | 0 | 0 | 3 |
| Aquatic | Trichoptera | Limnephilidae | Halesus sp. | 10 | 0 | 0 | 3 | 0 | 0 | 0 | 0 | 0 | 0 | 3 |
| Aquatic | Trichoptera | Limnephilidae | Micropterna sp. | 41 | 0 | 0 | 3 | 0 | 0 | 0 | 0 | 0 | 0 | 3 |
| Aquatic | Trichoptera | Limnephilidae | Potamophylax sp. | 188 | 0 | 0 | 3 | 1 | 0 | 0 | 0 | 0 | 0 | 3 |
| Aquatic | Trichoptera | Limnephilidae | Stenophylax sp. | 15 | 0 | 0 | 3 | 0 | 0 | 0 | 0 | 0 | 0 | 3 |
| Aquatic | Trichoptera | Philopotamidae | Philopotamus sp. | 1 | 0 | 0 | 0 | 2 | 3 | 0 | 1 | 0 | 0 | 3 |
| Aquatic | Trichoptera | Polycentropodidae | Plectrocnemia sp. | 14 | 0 | 0 | 0 | 0 | 1 | 0 | 3 | 0 | 0 | 3 |
| Aquatic | Trichoptera | Psychomyiidae | Tinodes sp. | 2 | 0 | 1 | 0 | 3 | 2 | 0 | 1 | 0 | 0 | 3 |
| Aquatic | Trichoptera | Rhyacophilidae | Hyporhyacophila sp. | 10 | 0 | 0 | 2 | 0 | 0 | 0 | 1 | 0 | 0 | 3 |
| Aquatic | Trichoptera | Rhyacophilidae | Rhyacophila sp. | 23 | 0 | 0 | 0 | 0 | 0 | 0 | 3 | 0 | 0 | 3 |
| Aquatic | Trichoptera | Sericostomatidae | Sericostoma sp. | 17 | 0 | 0 | 3 | 1 | 0 | 0 | 0 | 0 | 0 | 3 |
| Aquatic | Tricladida | Planariidae | Polycelis felina | 64 | 0 | 0 | 0 | 0 | 0 | 0 | 3 | 0 | 0 | 3 |
| Aquatic | Turbellaria | Planariidae | Dendrocoelum lacteum | 13 | 0 | 0 | 0 | 0 | 0 | 0 | 3 | 0 | 0 | 3 |
| Aquatic | Turbellaria | Planariidae | Dugesia sp. | 38 | 0 | 0 | 0 | 0 | 0 | 0 | 3 | 0 | 0 | 3 |
| Aquatic/ Terrestrial | Oligochaeta | | Oligochaeta | 448 | 1 | 3 | 0 | 1 | 0 | 0 | 0 | 0 | 0 | 3 |
| Terrestrial | Acarina | | Acarina sp. | 1 | 0 | 0 | 0 | 0 | 0 | 3 | 0 | 0 | 1 | 2 |
| Terrestrial | Aranea | Anyphaenidae | Anyphaenidae | 1 | 0 | 0 | 0 | 0 | 0 | 0 | 3 | 0 | 0 | 5 |
| Terrestrial | Aranea | Dictynidae | Dictynidae | 2 | 0 | 0 | 0 | 0 | 0 | 0 | 3 | 0 | 0 | 5 |
| Terrestrial | Aranea | Lycosidae | Pardosa sp. | 2 | 0 | 0 | 0 | 0 | 0 | 0 | 3 | 0 | 0 | 5 |
| Terrestrial | Aranea | Synaphridae | Synaphris sp. | 1 | NA | NA | NA | NA | NA | NA | NA | NA | NA | NA |
| Terrestrial | Aranea | Tetragnathidae | Pachygnatha sp. | 1 | 0 | 0 | 0 | 0 | 0 | 0 | 3 | 0 | 0 | 5 |
| Terrestrial | Aranea | Tetragnathidae | Tetragnatha sp. | 1 | 0 | 0 | 0 | 0 | 0 | 0 | 3 | 0 | 0 | 5 |
| Terrestrial | Aranea | Theridiidae | Theridiidae | 5 | 0 | 0 | 0 | 0 | 0 | 0 | 3 | 0 | 0 | 5 |
| Terrestrial | Aranea | Zoriidae | Zoriidae | 1 | 0 | 0 | 0 | 0 | 0 | 0 | 3 | 0 | 0 | 5 |
| Terrestrial | Coleoptera | Anobiidae | Stegobium sp. | 1 | NA | NA | NA | NA | NA | NA | NA | NA | NA | NA |
| Terrestrial | Coleoptera | Carabidae | Carabidae | 24 | 0 | 0 | 0 | 1 | 0 | 0 | 3 | 0 | 0 | 5 |
| Terrestrial | Coleoptera | Coccinellidae | Halyzia sp. | 2 | 0 | 0 | 0 | 0 | 0 | 0 | 3 | 0 | 0 | 5 |
| Terrestrial | Coleoptera | Coccinellidae | Myzia oblongoguttata | 2 | 0 | 0 | 0 | 0 | 0 | 0 | 3 | 0 | 0 | 5 |
| Terrestrial | Coleoptera | Coccinellidae | Sticholotidinae | 1 | NA | NA | NA | NA | NA | NA | NA | NA | NA | NA |
| Terrestrial | Coleoptera | Curculionidae | Curculionidae | 5 | NA | NA | NA | NA | NA | NA | NA | NA | NA | NA |
| Terrestrial | Coleoptera | Latridiidae | Corticariinae | 2 | 0 | 0 | 1 | 2 | 0 | 0 | 0 | 0 | 0 | 5 |
| Terrestrial | Coleoptera | Melyridae | Attalus sp. | 1 | NA | NA | NA | NA | NA | NA | NA | NA | NA | NA |
| Terrestrial | Coleoptera | Scarabaeidae | Melonthinae | 17 | 0 | 3 | 0 | 0 | 0 | 0 | 0 | 0 | 0 | 5 |
| Terrestrial | Coleoptera | Staphylinidae | Staphylinidae | 51 | 0 | 0 | 1 | 1 | 0 | 0 | 3 | 0 | 0 | 5 |
| Terrestrial | Collembola | Actaletidae | Actaletidae | 1 | NA | NA | NA | NA | NA | NA | NA | NA | NA | NA |
| Terrestrial | Collembola | Entomobryidae | Entomobryidae | 2 | 0 | 2 | 3 | 1 | 0 | 0 | 0 | 0 | 0 | 6; 7 |
| Terrestrial | Collembola | Isotomidae | Isotomidae | 32 | 0 | 0 | 0 | 3 | 0 | 0 | 0 | 0 | 0 | 6; 7 |
| Terrestrial | Collembola | Orchesellidae | Orchesella sp. | 9 | 0 | 2 | 3 | 1 | 0 | 0 | 0 | 0 | 0 | 6; 7 |

| | | | | | | | | | | | | | | |
|---|---|---|---|---|---|---|---|---|---|---|---|---|---|---|
| Terrestrial | Collembola | Thomoceridae | Thomoceridae | 5 | 0 | 2 | 3 | 1 | 0 | 0 | 0 | 0 | 0 | 6; 7 |
| Terrestrial | Deccapoda | Cambaridae | Orconectes limosus | 1 | 0 | 0 | 2 | 2 | 0 | 0 | 2 | 2 | 0 | 3 |
| Terrestrial | Diplopoda | Craspedosomatidae | Craspedosoma sp. | 2 | 0 | 0 | 3 | 1 | 0 | 0 | 0 | 0 | 0 | 5 |
| Terrestrial | Diplopoda | Julidae | Julidae | 2 | 0 | 0 | 3 | 1 | 0 | 0 | 0 | 0 | 0 | 5 |
| Terrestrial | Diplopoda | Polyxenidae | Polyxenus lagurus | 12 | 0 | 0 | 1 | 3 | 0 | 0 | 0 | 0 | 0 | 5 |
| Terrestrial | Diptera | Cecidomyiidae | Cecidomyiidae | 1 | NA | NA | NA | NA | NA | NA | NA | NA | NA | NA |
| Terrestrial | Diptera | Chloropidae | Chloropidae | 1 | NA | NA | NA | NA | NA | NA | NA | NA | NA | NA |
| Terrestrial | Gasteropoda | Enidae | Merdigera obscura | 1 | 0 | 0 | 3 | 1 | 0 | 0 | 0 | 0 | 0 | 8 |
| Terrestrial | Gasteropoda | Helicidae | Cepaea sp. | 1 | 0 | 0 | 3 | 0 | 0 | 0 | 0 | 0 | 0 | 5 |
| Terretrial | Gasteropoda | Zonitidae | Zonitidae | 1 | 0 | 2 | 2 | 2 | 0 | 0 | 2 | 0 | 0 | 3 |
| Terrestrial | Gasteropoda | Valloniidae | Acanthinula aculeata | 1 | NA | NA | NA | NA | NA | NA | NA | NA | NA | NA |
| Terrestrial | Geophilomorpha | Geophilidae | Geophilus pusillifrater | 1 | 0 | 0 | 0 | 0 | 0 | 0 | 3 | 0 | 0 | 9 |
| Terrestrial | Geophilomorpha | Linotaeniidae | Strigamia sp. | 1 | 0 | 0 | 0 | 0 | 0 | 0 | 3 | 0 | 0 | 9 |
| Terrestrial | Hemiptera | Diaspididae | Diaspididae | 1 | 0 | 0 | 0 | 0 | 0 | 3 | 0 | 0 | 0 | 5 |
| Terrestrial | Hemiptera | Hydrometridae | Hydrometra sp. | 2 | 0 | 0 | 0 | 0 | 0 | 3 | 0 | 0 | 0 | 5 |
| Terrestrial | Hemiptera | Miridae | Miridae | 1 | 0 | 0 | 0 | 0 | 0 | 3 | 0 | 0 | 0 | 5 |
| Terrestrial | Hymenoptera | Formicidae | Lasius sp. | 3 | NA | NA | NA | NA | NA | NA | NA | NA | NA | NA |
| Terrestrial | Hymenoptera | Formicidae | Myrmica rubra | 3 | NA | NA | NA | NA | NA | NA | NA | NA | NA | NA |
| Terrestrial | Hymenoptera | Formicidae | Temnothorax sp. | 1 | NA | NA | NA | NA | NA | NA | NA | NA | NA | NA |
| Terrestrial | Isopoda | Armadillidiidae | Armadillidium sp. | 26 | 0 | 0 | 3 | 0 | 0 | 0 | 0 | 0 | 0 | 5 |
| Terrestrial | Isopoda | Cylisticidae | Cylisticus sp. | 3 | 0 | 0 | 3 | 0 | 0 | 0 | 0 | 0 | 0 | 5 |
| Terrestrial | Isopoda | Oniscidae | Oniscus asellus | 1 | 0 | 0 | 3 | 0 | 0 | 0 | 0 | 0 | 0 | 5 |
| Terrestrial | Isopoda | Trichoniscidae | Trichoniscus sp. | 7 | 0 | 0 | 3 | 0 | 0 | 0 | 0 | 0 | 0 | 5 |
| Terrestrial | Isopoda | Philosciidae | Philoscia sp. | 14 | 0 | 0 | 3 | 0 | 0 | 0 | 0 | 0 | 0 | 5 |
| Terrestrial | Isopoda | Porcellionidae | Porcellionidae | 21 | 0 | 0 | 3 | 0 | 0 | 0 | 0 | 0 | 0 | 5 |
| Terrestrial | Lepidoptera | Noctuidae | Noctuidae | 1 | 0 | 0 | 2 | 2 | 0 | 0 | 0 | 0 | 0 | 5 |
| Terrestrial | Lithobiomorpha | Lithobiidae | Lithobius sp. | 14 | 0 | 0 | 1 | 0 | 0 | 0 | 3 | 0 | 0 | 10 |
| Terrestrial | Odonata | Calopterygidae | Calopteryx sp. | 5 | 0 | 0 | 0 | 0 | 0 | 0 | 3 | 0 | 0 | 5 |
| Terrestrial | Pseudoscorpiones | Chthoniidae | Chthoniidae | 1 | NA | NA | NA | NA | NA | NA | NA | NA | NA | NA |
| Terrestrial | Psocoptera | Ectopsocidae | Ectopsocus sp. | 2 | NA | NA | NA | NA | NA | NA | NA | NA | NA | NA |


1. Schmidt-Kloiber, A. & Hering, D. (2015) www.freshwaterecology.info – An online tool that unifies, standardises and codifies more than 20,000 European freshwater organisms and their ecological preferences. Ecological Indicators, 53, 271-282.
2. Di Sabatino, A., Gerecke, R. and Martin, P. (2000), The biology and ecology of lotic water mites (Hydrachnidia). Freshwater Biology, 44: 47-62. https://doi.org/10.1046/j.1365-2427.2000.00591.x
3. Tachet, H., Richoux, P., Bournaud, M. & Usseglio-Polatera, P. (2010) Invertébrés d'Eau Douce (3rd edition)., CNRS. Paris.
4. Nentwig, W. The prey of web-building spiders compared with feeding experiments (Araneae: Araneidae, Linyphiidae, pholcidae, Agelenidae). *Oecologia* **56**, 132–139 (1983). https://doi.org/10.1007/BF00378229
5. Webb, J., Heaver, D., Lott, D., Dean, H.J., van Breda, J., Curson, J., Harvey, M.C., Gurney, M., Roy, D.B., van Breda, A., Drake, M., Alexander, K.N.A. and Foster, G. (2018). Pantheon - database version 3.7.6
6. Berg, M, P., Stoffer, M. and van den Heuvel, H, H. 2004 Feeding guilds in Collembola based on digestive enzymes. Pedobiologia 48, 589-601. https://doi.org/10.1016/j.pedobi.2004.07.006.

7. Chahartaghi, M., Langel., R., Scheu, S. and Ruess, L. (2005) Feeding guilds in Collembola based on nitrogen stable isotope ratios. Soil Biology and Biochemistry 37, 1718-1725 https://doi.org/10.1016/j.soilbio.2005.02.006.
8. Wäreborn, I. (1969). Land Molluscs and Their Environments in an Oligotrophic Area in Southern Sweden. *Oikos*, *20*(2), 461–479. https://doi.org/10.2307/3543209
9. Bortolin, F., Fusco, G., Bonato, L. 2018. Comparative analysis of diet in syntopic geophilomorph species (Chilopoda, Geophilomorpha) using a DNA-based approach. Soil Biology and Biochemistry 127, 223-229. https://doi.org/10.1016/j.soilbio.2018.09.021.
10. Lewis, J, G, E. (1965) The food and reproductive cycles of the centipedes *Lithobius variegatus* and *Lithobius forficatus* in a Yorkshire woodland. Proceedings of the Zoological Society of London, 144: 269-284. https://doi.org/10.1111/j.1469-7998.1965.tb05178.x

**Table S2.** Outputs from the linear model testing for the effects of flow intermittence (FI), upstream distance to source (DS) and upstream proportion of perennial reaches (UC) and their interactions on four attributes of resilience, i.e. the number of scales (Nb of scales), within-scale richness, scale span and gap size. Significant effects ($P < 0.05$) are indicated in bold and near-significant ($0.05 < P < 0.10$) are in italics.

| | **Nb of scales** | | **Within-scale richness** | | **Scale span** | | **Gap size** | |
|---|---|---|---|---|---|---|---|---|
| | F | P | F | P | F | P | F | P |
| FI | 15,01 | **0,002** | 6,90 | **0,02** | 1,05 | 0,33 | 0,19 | 0,67 |
| DS | 0,97 | 0,32 | 5,36 | **0,04** | 2,98 | 0,11 | 1,38 | 0,26 |
| UC | 0,55 | 0,43 | 0,18 | 0,68 | 2,06 | 0,18 | 0,33 | 0,58 |
| FI*DS | 4,68 | **0,05** | 1,29 | 0,28 | 0,09 | 0,77 | 10,89 | **0,006** |
| FI*UC | 0,30 | 0,59 | 0,96 | 0,35 | 0,14 | 0,71 | 0,39 | 0,54 |
| DS*UC | 5,99 | **0,03** | 0,82 | 0,38 | 6,08 | **0,03** | 1,17 | 0,30 |
| FI*DS*UC | 0,05 | 0,83 | 3,23 | *0,10* | 0,18 | 0,68 | 5,28 | **0,04** |

**Table S3.** Outputs from the linear model testing for the effects of flow intermittence (FI), upstream distance to source (DS) and upstream proportion of perennial reaches (UC) and their interactions on cross-scale functional redundancy in different functional feeding groups such as deposit-feeders, filter-feeders, grazers, insectivores, parasites, piercers, piscivores and shredders. Significant effects (P < 0.05) are indicated in bold and near-significant (0.05 < P < 0.10) are in italics.

| | Deposit-feeders | | Filter-feeders | | Grazers | | Insectivores | | Parasites | | Piercers | | Piscivores | | Shredders | |
|---|---|---|---|---|---|---|---|---|---|---|---|---|---|---|---|---|
| | F | P | F | P | F | P | F | P | F | P | F | P | F | P | F | P |
| FI | 1,12 | 0,31 | 5,54 | **0,04** | 9,97 | **0,008** | 14,79 | **0,002** | 0,51 | 0,49 | 4,50 | **0,05** | 0,06 | 0,82 | 4,85 | **0,05** |
| DS | 0,0001 | 0,99 | 2,73 | 0,12 | 2,67 | 0,13 | 13,82 | **0,003** | 1,56 | 0,24 | 5,94 | **0,03** | 0,89 | 0,37 | 2,05 | 0,18 |
| UC | 0,20 | 0,66 | 0,18 | 0,68 | 0,03 | 0,86 | 1,47 | 0,25 | 0,40 | 0,54 | 0,09 | 0,77 | 0,12 | 0,74 | 5,42 | **0,04** |
| FI*DS | 0,02 | 0,88 | 6,92 | **0,02** | 2,68 | 0,13 | 11,12 | **0,006** | 0,82 | 0,38 | 4,11 | *0,07* | 0,25 | 0,63 | 0,31 | 0,59 |
| FI*UC | 0,002 | 0,97 | 2,73 | 0,12 | 1,96 | 0,19 | 0,07 | 0,79 | 0,003 | 0,96 | 2,59 | 0,13 | 0,005 | 0,94 | 0,43 | 0,53 |
| DS*UC | 4,32 | *0,06* | 2,51 | 0,14 | 4,16 | *0,06* | 5,28 | **0,04** | 1,40 | 0,26 | 0,003 | 0,96 | 1,53 | 0,24 | 4,41 | *0,06* |
| FI*DS*UC | 2,27 | 0,16 | 0,77 | 0,40 | 0,13 | 0,72 | 0,04 | 0,85 | 0,14 | 0,71 | 0,29 | 0,60 | 0,07 | 0,79 | 0,06 | 0,81 |

**Table S4.** Outputs from the linear model testing for the effects of flow intermittence (FI), upstream distance to source (DS) and upstream proportion of perennial reaches (UC) and their interactions on within-scale functional redundancy in different functional feeding groups such as deposit-feeders, filter-feeders, grazers, insectivores, parasites, piercers and shredders. Models for within-scale functional redundancy in piscivores could not be computed as within-scale functional redundancy in piscivores was null for all stream sites, except in one intermittent stream. Significant effects (P < 0.05) are indicated in bold and near-significant (0.05 < P < 0.10) are in italics.

| | **Deposit-feeders** | | **Filter-feeders** | | **Grazers** | | **Insectivores** | | **Parasites** | | **Piercers** | | **Piscivores** | | **Shredders** | |
|---|---|---|---|---|---|---|---|---|---|---|---|---|---|---|---|---|
| | F | P | F | P | F | P | F | P | F | P | F | P | F | P | F | P |
| FP | 0,35 | 0,56 | 2,76 | 0,12 | 13,73 | **0,003** | 1,8 | 0,20 | 0,51 | 0,49 | 0,82 | 0,38 | - | - | 0,99 | 0,34 |
| DS | 0,02 | 0,9 | 2,24 | 0,16 | 3,34 | *0,09* | 13,5 | **0,003** | 1,56 | 0,24 | 7,56 | **0,02** | - | - | 0,42 | 0,53 |
| UC | 1,68 | 0,22 | 0,004 | 0,95 | 1,69 | 0,22 | 2,43 | 0,15 | 0,40 | 0,54 | 0,001 | 0,98 | - | - | 0,93 | 0,35 |
| FP*DS | 2,69 | 0,13 | 8,55 | **0,01** | 0,80 | 0,39 | 6,27 | **0,03** | 0,82 | 0,38 | 2,09 | 0,17 | - | - | 0,004 | 0,95 |
| FP*UC | 0,19 | 0,67 | 3,02 | 0,11 | 1,49 | 0,25 | 0,73 | 0,41 | 0,003 | 0,96 | 2,20 | 0,16 | - | - | 0,23 | 0,64 |
| DS*UC | 2,90 | 0,11 | 3,28 | *0,10* | 0,07 | 0,80 | 0,59 | 0,46 | 1,40 | 0,26 | 0,00 | 0,99 | - | - | 0,50 | 0,49 |
| FP*DS*UC | 3,78 | *0,08* | 0,38 | 0,55 | 1,19 | 0,30 | 0,007 | 0,93 | 0,14 | 0,71 | 0,37 | 0,55 | - | - | 1,05 | 0,33 |

**Table S5.** Outputs from the linear model testing for the effects of flow intermittence (FI), upstream distance to source (DS) and upstream proportion of perennial reaches (UC) and their interactions on response diversity of different functional feeding groups such as deposit-feeders, filter-feeders, grazers, insectivores, parasites, piercers, piscivores and shredders. Significant effects (P < 0.05) are indicated in bold and near-significant (0.05 < P < 0.10) are in italics.

| | **Deposit-feeders** | | **Grazers** | | **Insectivores** | | **Piercers** | | **Shredders** | |
|---|---|---|---|---|---|---|---|---|---|---|
| | F | P | F | P | F | P | F | P | F | P |
| FI | 2,47 | 0,14 | 0,85 | 0,38 | 1,5 | 0,24 | 1,9 | 0,22 | 14,63 | **0,002** |
| DS | 1,51 | 0,24 | 0,11 | 0,75 | 0,23 | 0,64 | 0,56 | 0,48 | 4,95 | **0,05** |
| UC | 0,0003 | 0,99 | 4,65 | **0,05** | 1,02 | 0,33 | 2,41 | 0,17 | 0,82 | 0,38 |
| FI*DS | 0,31 | 0,59 | 1,19 | 0,30 | 0,68 | 0,42 | 5,1 | *0,06* | 1,08 | 0,32 |
| FI*UC | 0,42 | 0,53 | 0,87 | 0,37 | 0,02 | 0,88 | 11,1 | **0,02** | 1,63 | 0,23 |
| DS*UC | 0,01 | 0,94 | 0,96 | 0,35 | 0,18 | 0,68 | 0,21 | 0,67 | 4,35 | *0,06* |
| FI*DS*UC | 0,0007 | 0,98 | 4,47 | **0,05** | 0,26 | 0,62 | 0,13 | 0,73 | 0,86 | 0,37 |

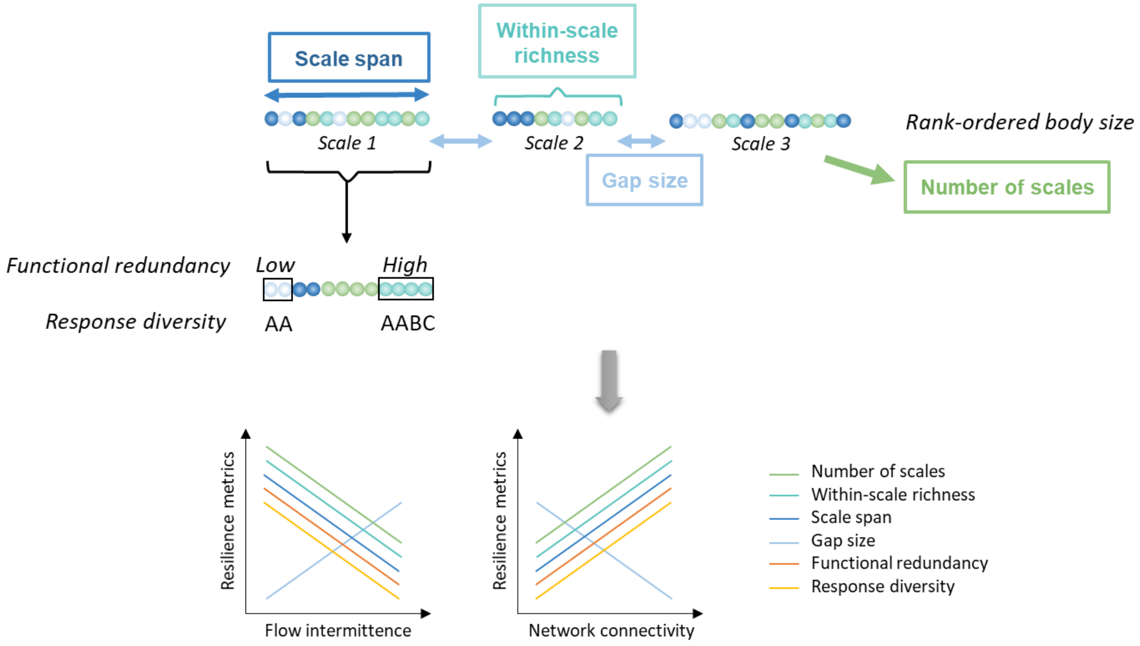


**Figure S1**. Conceptual figure showing the different metrics assessed within the cross-scale model of resilience (upper part of the figure) and the expected relationships between these metrics and flow intermittence and network connectivity.

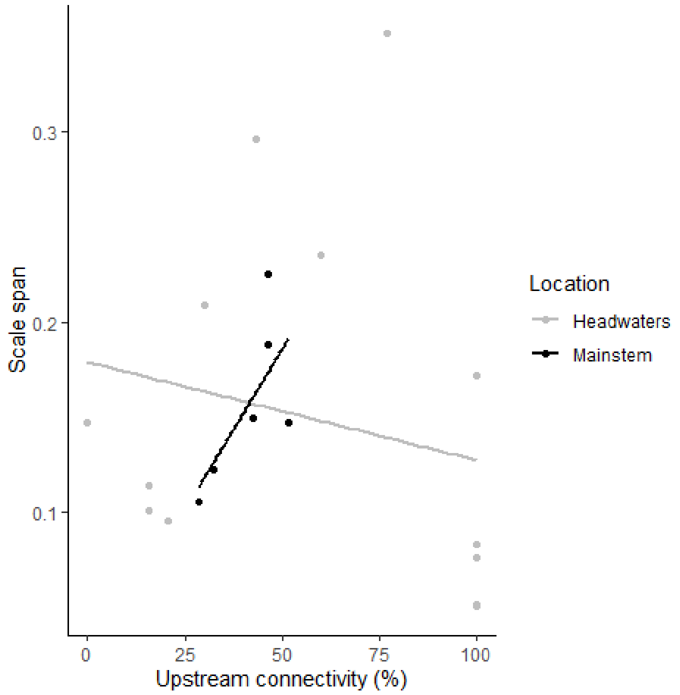


**Figure S2.** Response of scale span to upstream proportion of perennial reaches (upstream connectivity). Scale span is defined as the average difference between the greatest and smallest logtransformed body size within the aggregation. These aggregation groups were identified by the discontinuity analysis applied to body size distribution.

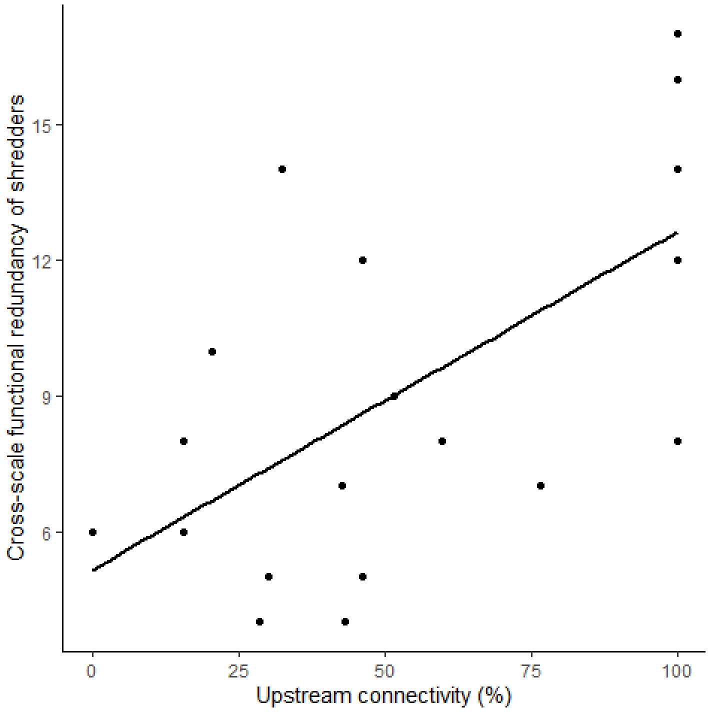


**Figure S3.** Responses of the cross-scale functional redundancy of shredders to upstream proportion of perennial reaches (upstream connectivity). Cross-scale functional redundancy is defined as the number of functional feeding groups present at each scale (i.e. body size aggregation).

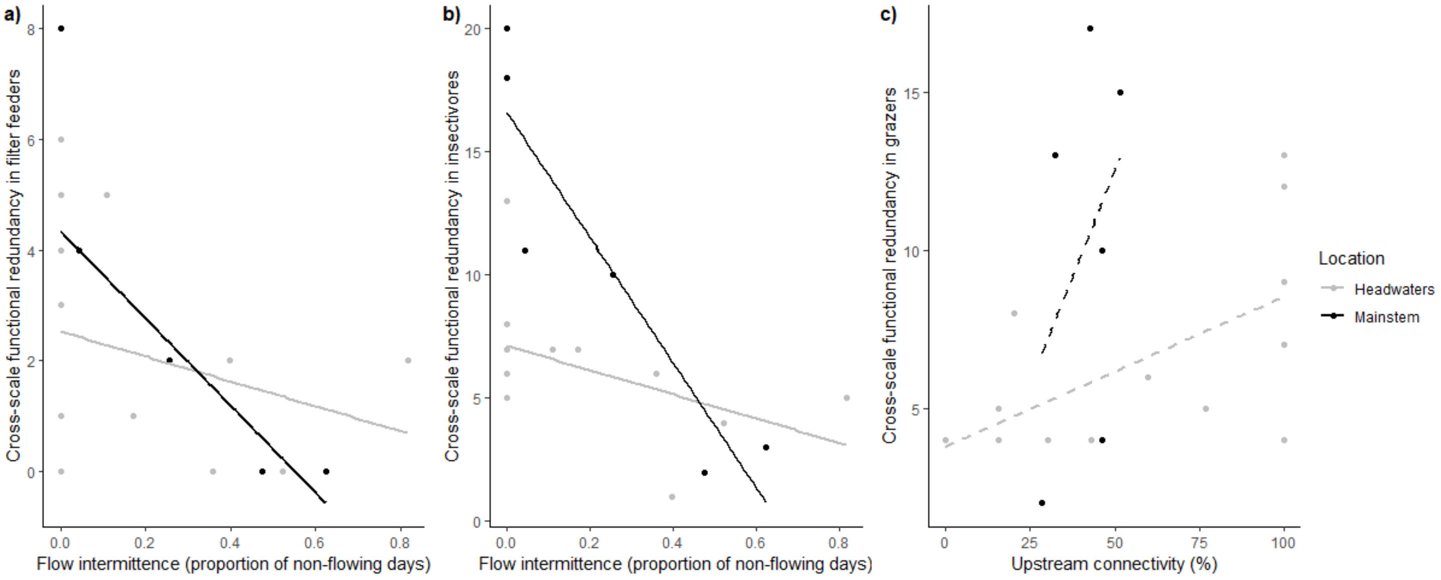


**Figure S4.** Response of cross-scale functional redundancy in filter-feeders (a), insectivores (b) and grazers (c) to flow intermittence (a, b) and upstream proportion of perennial reaches (upstream connectivity; c), depending on site location (headwaters vs. mainstem).

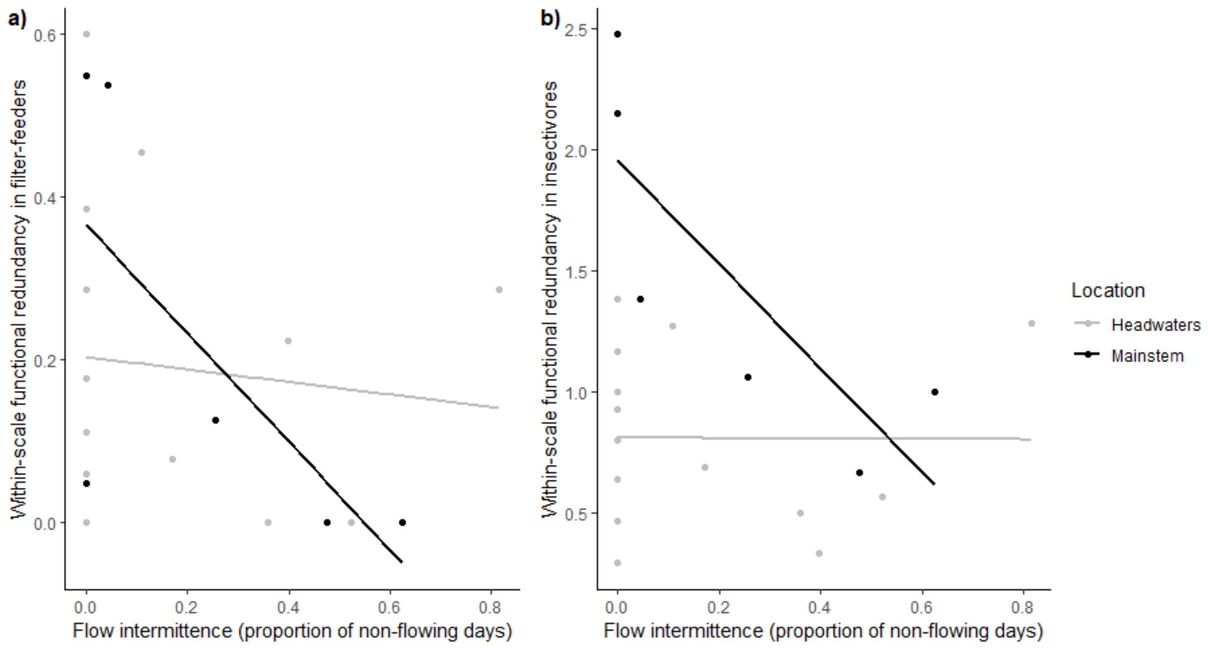


**Figure S5.** Response of within-scale functional redundancy in filter-feeders (a) an insectivores (b) to flow intermittence, depending on site location (headwaters vs. mainstem).

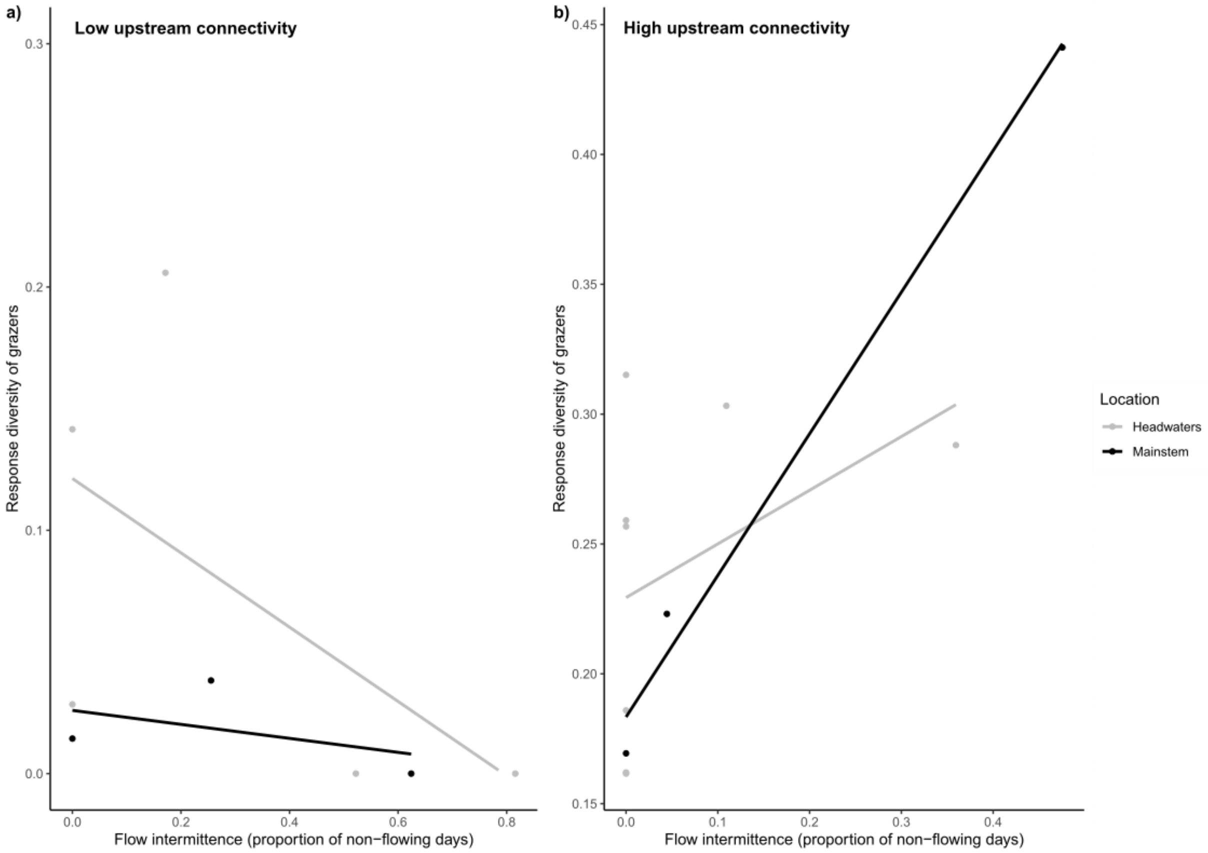


**Figure S6.** Response diversity of grazers to flow intermittence, depending on site location (headwaters vs. mainstem) under low upstream connectivity (a) and high upstream connectivity (b).

*Supplementary material to Truchy, Sarremejane, Braun, Datry. Community resilience in a river network: the roles of connectivity and drying regime*
